# Temperature moderates eDNA-biomass relationships in northern pike

**DOI:** 10.1101/2022.12.28.522080

**Authors:** M. Ogonowski, E. Karlsson, A. Vasemägi, J. Sundin, P. Bohman, G. Sundblad

**Affiliations:** Department of Aquatic Resources, Institute of Freshwater Research, Swedish University of Agricultural Sciences, Stångholmsvägen 2, SE-17893, Drottningholm, Sweden; Chair of Aquaculture, Institute of Veterinary Medicine and Animal Sciences, Estonian University of Life Sciences, Kreutzwaldi 46, 51006, Tartu, Estonia

**Keywords:** *Esox lucius*, angling, CPUE, eDNA, environmental DNA, biomass, abundance, temperature, spawning, coast, Baltic Sea

## Abstract

Support for eDNA as a quantitative monitoring tool is growing worldwide. Despite advances there are still uncertainties regarding the representability of the eDNA signal over varying spatiotemporal scales, influence of abiotic forcing and phenological changes affecting behavior of the study organism, particularly in open environments. To assess the spatiotemporal variability and predictive power of quantitative eDNA analysis, we applied species-specific real-time quantitative PCR on water filtrates during two visits to 22 coastal bays in the Baltic Sea. Within bays, we collected water along four transects across each bay and compared the pooled eDNA concentration to temporally matched catches from standardized angling targeting the northern pike (*Esox lucius*) a species for which reliable monitoring data is lacking. We found the variability in eDNA concentrations between transects to be moderate (21%) but still considerably lower than across bays and visits (52%), suggesting small scale spatial differences are of less importance during spring when pike spawn. Standardized angling catches, bay area, and water temperature together explained 48% of the variance in eDNA concentrations. DNA concentrations decreased with increasing bay area, likely indicating a dilution effect. Notably, the relationship between eDNA and standardized catches was positive but varied with temperature and the eDNA-abundance relationship was only significant at higher temperatures, which also coincided with a higher proportion of spawning/spent fish. We conclude that temperature is a key moderating factor driving changes in pike behaviour and spring DNA-dynamics. We recommend that future surveys focus on larger spatiotemporal scales during times when the influence of changing temperatures is minimized.

## 1. Introduction

The monitoring of fish stocks requires quantitative data on their abundance. This can be difficult to obtain for species with sedentary lifestyles and whose catchability in passive gears, such as gillnets and traps, is restricted (Villegas-Ríos et al. 2014). Environmental DNA (eDNA) has been suggested as a possible tool for fish stock monitoring in general, and monitoring of species with low catchability in particular (Kačergytė et al. 2021). While eDNA can be successfully used in biodiversity monitoring based on presence/absence data (Dejean et al. 2011, Thomsen et al. 2012, Takahara et al. 2013, Dunker et al. 2016, Hernandez et al. 2020), it’s potential use for biomass estimation is still under development. Quantitative relationships between eDNA concentrations and fish biomass have been demonstrated under controlled conditions with known biomass and to a lesser extent also in natural environments with unknown biomass (Rourke et al. 2021 and references therein). Although linear relationships have been obtained in controlled experiments, the precision of these estimates varies greatly in natural systems where eDNA on average explains 57% of the variance compared to 81 % in controlled mesocosm experiments (Yates et al. 2019). The variability in these estimates can be attributed to differences in ground truthing methods (Rourke et al. 2021), ambient temperature (Lacoursière-Roussel et al. 2016), hydrologic conditions (Song et al. 2017), DNA extraction methods (Bockrath et al. 2022, Karlsson et al. 2022), presence of Polymerase Chain Reaction (PCR) inhibiting humic and tannic acids (Lance and Guan 2020), sediment particles in the water (Stoeckle et al. 2017) and, size distribution of the local population (Yates et al. 2021a, 2021c). Moreover, the positive relationship between eDNA and fish biomass in the wild has most often been found in smaller enclosed lake systems (Spear et al. 2021), streams (Yates et al. 2021a) and to some extent pelagic marine environments (Li et al. 2022). Comparative eDNA surveys for semi-open coastal fish communities are still scarce in the literature (Rourke et al. 2021).

The northern pike (*Esox lucius*, Linnaeus, 1758) is a species of growing research interest (Forsman et al. 2015). It is a keystone predator in freshwater and coastal ecosystems and it is important for ecosystem functioning as well as a focal species for the recreational fishery. Pike also provides an example of a species for which accurate abundance indices are difficult to obtain due to its low catchability in passive gears (Craig 2008). In the Baltic Sea, large scale patterns indicate that the pike populations on the east coast of Sweden have drastically declined. The reasons are multifaceted but likely a consequence of increased predation on adults from grey seals and cormorants (Hansson et al. 2017, Svensson 2021), predation on juvenile stages by three-spined stickleback (Eklöf et al. 2020, Donadi et al. 2020), loss of recruitment habitats (Sundblad and Bergström 2014), and also a period of high recreational fishing mortality during the early 1990s (Bergström et al. 2022). For stationary species which form genetically stable distinct populations over rather small geographical areas (Laikre et al. 2005, Wennerström et al. 2016, Möller et al. 2021, Diaz-Suarez et al. 2022) management needs to be regional and there is a need for monitoring methods which are able to accurately assess the status of pike populations on a local scale. Northern pike aggregate in shallow areas to spawn during spring and have a strong homing behavior. This requires monitoring with a high level of spatial coverage, which poses challenges to the management of this species. Since traditional, passive and lethal monitoring methods have proven ineffective, recent attempts to quantify pike abundance have employed active methods such as standardized rod-and-reel fishing during the spawning period to obtain measures of relative abundance (Catch-Per-Unit-Effort data, CPUE) and size structure of distinct populations that form local spawning aggregations. Standardization of such methods is however complicated as the size and type of bait used, catch-and-release (C&R) practices and angling effort can affect the catchability (Arlinghaus et al. 2008, 2017, Kuparinen et al. 2010), meaning that CPUE can be underestimated in areas where fishing is intense and C&R is common. eDNA on the other hand offers several advantages over active rod fishing in the sense that it is not size selective, unaffected by fishing effort and gear use, can provide an adequate level of replication (Shelton et al. 2022), is cost-efficient and, since it is non-invasive, it potentially has a higher probability of better reflecting the local density of fish (Wilcox et al. 2016).

Strong positive relationships between eDNA and biomass of pike have been shown in large outdoor mesocosms during the reproductive period (Karlsson et al. 2022). However, it is unknown if eDNA can provide quantitative data on pike abundance also under natural conditions. In this paper, we test the hypothesis that relative pike population sizes can be estimated using eDNA during the reproductive season when pike aggregate. We do this using a large number of coastal bays in the Baltic Sea where we compare eDNA concentrations to standardized angling. The aim is to investigate whether eDNA can be used as a non-invasive tool to measure fish abundance/biomass in semi-open coastal areas.

## 2. Materials & Methods

### 2.1 General design

To assess the potential of eDNA to estimate pike population biomass/abundance under natural conditions, we collaborated with a project conducting standardized angling to support management actions. The multi-year project was initiated by the Stockholm County Administrative Board and aimed to assess pike population sizes in relation to (current and future) fishing closures during the pike spawning season.

The study area covered >200 km of the Stockholm archipelago in the Baltic Sea. The angling was performed during two visits in 24 coastal bays during April-May 2020 (coordinates for each bay can be found in the supplementary data file “DATA.xlsx”). The selection of bays to include in this study was therefore reliant on the evaluation of fishing closures. The design for that evaluation was based on paired bays, of which some used a BACI-design. The paired bays were chosen to be in close proximity to each other and to be as similar as possible in terms of size, mean depth and habitat conditions, but with one bay being either closed or soon-to be closed for fishing and the other one open to angling; thus likely providing a range of fish densities spanning from low to high.

The eDNA sampling was performed in 22 out of the 24 fished bays a few days prior to each angling visit in a bay, in order to not risk the eDNA signal to be influenced by the fishing activity nor to disturb the fishing by simultaneously sampling eDNA (Figure 1).

**Figure 1.**
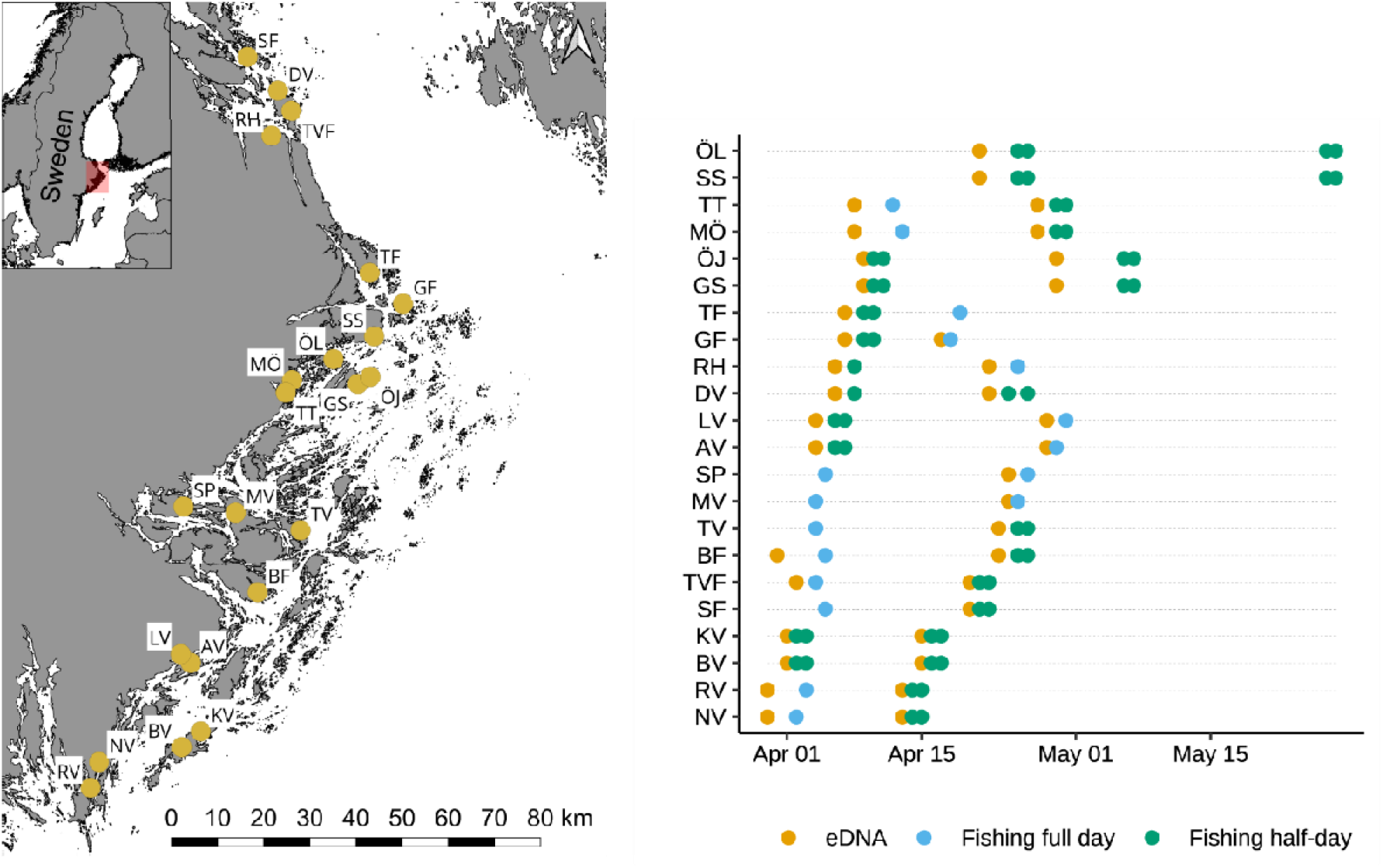
Overview map of bays sampled for eDNA and pike by angling (left panel) and sampling scheme (right panel) showing sampling dates for eDNA and angling ordered by bay-pair during the survey in 2020. Angling was either divided into two half-days within a bay-pair and fished for two consecutive days, alternating morning and afternoon in each bay, or as a full day’s fishing in a specific bay.

### 2.2 DNA analyses

#### 2.2.1 eDNA collection and filtration

Within each bay and visit, we collected water along four transects: three shallow water transects (A, B and C), each trailing roughly a third of the coastal length of the bay, and one deep water transect (D) across the center of the bay (Figure 2, supplementary list of figures “Bay.info.pdf”). For each transect, one liter of surface water was collected every 50 m. The distance between individual subsamples was chosen based on the reported detection distance for caged northern pike carcasses in a freshwater system (Dunker et al. 2016) and live Japanese striped jack (*Pseudocaranx dentex*, Bloch & Schneider, 1801) in a marine setting (Murakami et al. 2019). The total amount of water collected per bay and transect was approximately proportional to the bay area. The water from each transect was pooled in a large plastic container and the total volume of pooled water varied from 4 to 26 L, median = 10, IQR = 7 (Supporting information file, “DATA.xlsx”), depending on the length of the transect. From this pool of water, duplicate samples of one liter each were filtered using an established filtration technique (Karlsson et al. 2022) with some modifications. Water was pushed through a Swinnex filter holder (Merck KGaA, Darmstadt, Germany) loaded with two filters (cellulose nitrate filter, pore size of 0.8 μm and a glass microfiber filter (GFFA, pore size of approximately 1.6 μm, GE Healthcare, Chicago, United States)) using a plastic syringe. The glass microfiber filter functioned as a pre-filter that allowed a larger volume of water to pass through (Capo et al. 2020). We re-used the filter holder and exchanged the filters in the field for the second technical replicate. Although some cross-contamination could be expected at this stage, we assumed the contamination would be diluted to the point that it would fall below the detection limit. This was later confirmed by finding no consistent increase in eDNA concentrations in consecutive eDNA samples (Figure S 1). One field negative control using one liter of distilled water was run per bay visit directly after the filtration of the eDNA samples. After filtration, the filters were enclosed in zip-loc bags and snap-frozen on dry ice until arrival to the laboratory where they were directly transferred to a −80 °C freezer pending DNA extraction. Nitrile gloves and sterile pincers were used at all times during filter handling. All equipment was sterilized between field visits by immersion in 10%–20% commercial grade bleach for a minimum of 10 min and then rinsed thoroughly in tap water.

**Figure 2.**
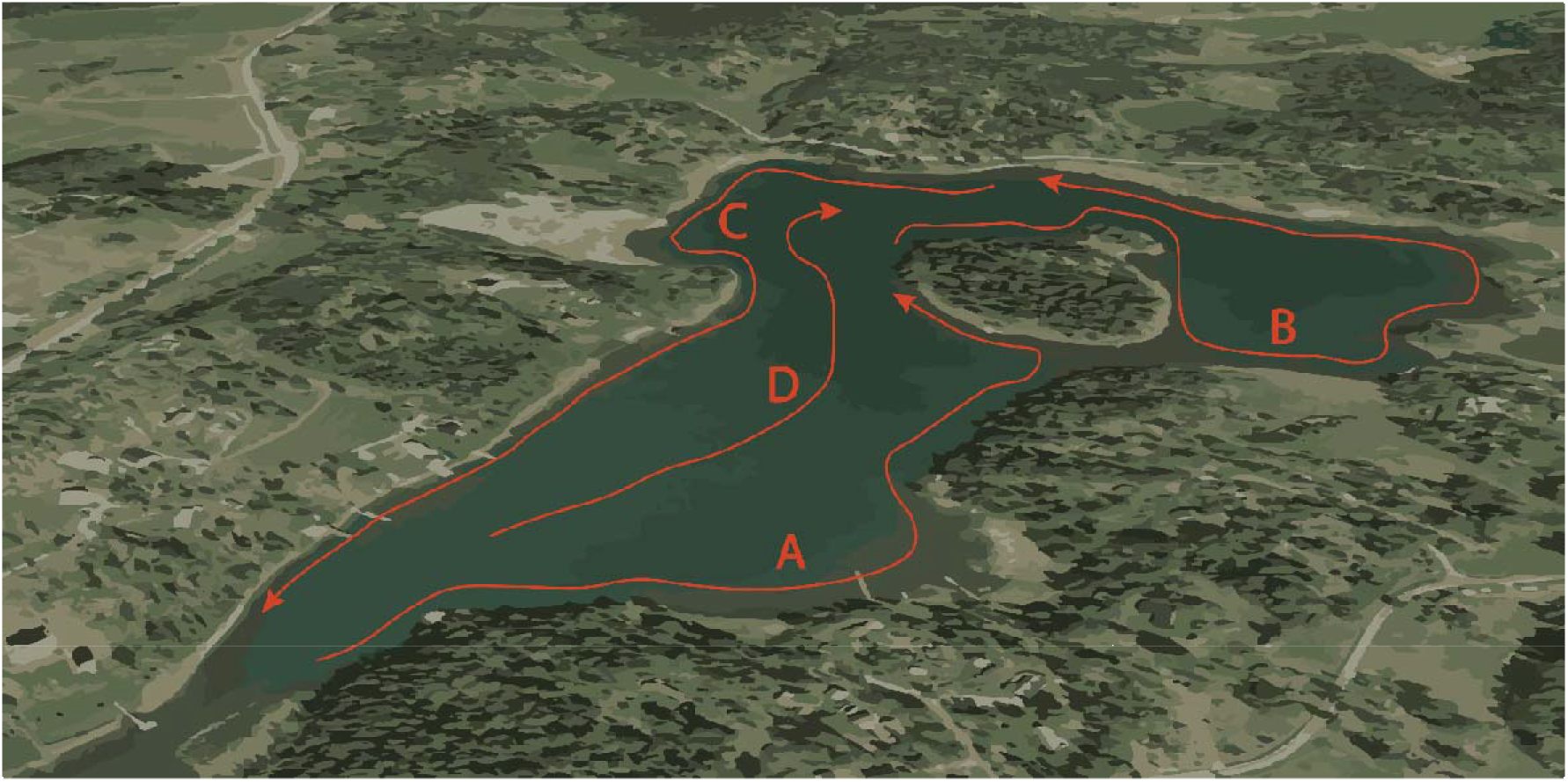
Schematic image showing the sampling design for eDNA. 1 L water samples were collected 50 m apart in four transects (A-D) and pooled within each transect. Transects A-C normally covered the shallowest vegetated areas while transect D was the deepest in the central part of the bay. Transect length was approximately proportional to the bay area.

#### 2.2.2 DNA extraction

DNA was extracted from the filters using a modified Chelex extraction protocol described in Karlsson et al. (2022). In brief, the filters were cut into smaller pieces using sterile equipment and then mixed with 750 μl of a 10% (w/v) Chelex suspension in 5 mL Eppendorf ® screw cap tubes. The tubes containing the filter cuttings were heated at 100 °C for 10 min to lyse cell material and denaturate the DNA, and then vortexed thoroughly. This step was repeated twice after which the supernatant was transferred to a smaller 1.5 mL tube and centrifuged at 16 × 10^3^ g for 1.5 min to remove remaining filter debris and Chelex from the solution. After centrifugation, the supernatant was once again transferred to a clean 1.5 mL tube. If necessary, any remaining Chelex was removed by repeating the last centrifugation step and transferring the supernatant to a clean tube.

#### 2.2.3 DNA quantification using qPCR

We used a real-time quantitative polymerase reaction assay (qPCR) for quantification of pike DNA in collected samples. The primer and probe combination (F-primer: 5’-CCT TCCCCCGCATAAATAATATAA-3’, R-primer: 5’-GTACCAGCACCAGCTTCAACAC-3’ and probe: 5’-FAM-CTTCTG ACTTCTCCCC-BHQ-1-3’ (Microsynth AG)) was originally developed by Olsen et al. (2015, 2016) and has later been successfully used for pike detection in water samples (Dunker et al. 2016, Karlsson et al. 2022). The assay targets a 94-base-pair-long fragment of the Cytochrome oxidase I gene (COI). qPCR was performed on a BioRad CFX364 Real-time PCR system with 15 μl reaction volumes. Reaction concentrations of the forward primer, reverse primer, and probe were 900 nM each with 7.5 μl 2× TaqMan™ Environmental Master Mix 2.0 (Thermo Fisher) in each well loaded with 4 μl of the sample template. An internal positive control (IPC) (Cy®5-QXL®670 Probe, EuroGentec, Liege, Belgium) kit was run in duplex reactions to control for potential inhibition. 0.3 μl of 10 × IPC mix and 0.2 μl of IPC template DNA was added to each reaction.

Inhibition in eDNA samples was determined based on aberrant IPC Cq-values. The expected Cq over the range of the standard curve was on average 27.3 Cq with average minimum and maximum values ranging 26.5–28.5 Cq. Therefore, we classified samples > 28.5 Cq as unacceptable. Such samples were purified using a Zymo OneStep PCR Inhibitor Removal Kit and reanalysed in the qPCR. If purification of the sample did not improve the IPC-value to within acceptable Cq-limits it was excluded (Figure S 2).

The following qPCR program was used for all the reactions: 10 min activation at 95 °C followed by 45 cycles of 15 s at 95 °C and 60 s at 60 °C. eDNA quantification was achieved using a standard curve consisting of an 8-step, 10-fold dilution series of pike DNA (1 – 1 × 10^7^ copies μl^-1^) with the addition of a lowest concentration at 0.25 copies μl^-1^. As a standard we used a synthetic 94 nucleotide oligo template targeting the mitochondrial COI-gene: 5’-CCT TCC CCC GCA TAA ATA ATA TAAGCT TCT GAC TTCTCCCCC CCT CCT TTT TAC TTC TCT TAG CCT CCT CAG TTC TCT GTG TTG AAG CTG GTG CTG GTA C-3’ and complementary strand: 5’-GTA CCA GCA CCA GCT TCA ACA CCT GAG GAG GCT AAG AGA AGT AAA AAG GAG GGG GGG AGA AGT CAG AAG CTT ATA TTA TTT ATG CGG GGG AAG G-3’ (Microsynth AG).

Samples and standard curves were run in quadruplicates with four no template control (NTC) reactions on each plate. Plate efficiency varied between 101.3% and 110.7%, with R^2^ values between 0.983 and 0.995.

#### 2.2.4 Determination of limits of detection (LOD) and quantification (LOQ)

Limit of detection (LOD) and limit of quantification (LOQ) were determined by running a standard curve with DNA concentrations in the same range as all other standards ranging 0.25–1 × 10^7^ copies μl^-1^ each in 16 technical replicates. LOD is defined as the lowest concentration of DNA where 95% of the technical replicates amplify and LOQ is defined as the lowest concentration of DNA with a coefficient of variation (CV) below 35% (Klymus et al. 2020). Effective LOD is defined as the lowest concentration with a detection probability of 95% given *n* replicates. The estimated qPCR efficiency was 118.2 % with R^2^ = 0.981. LOD and LOQ were both determined to 1.97 copies μl^-1^. Analysis in quadruplicates gave an effective LOD of 0.58 copies μl^-1^. Concentrations are given per microliter of target sample (4 μl).

#### 2.3.5. qPCR data handling and curation

Values below the LOD were set to the LOD while values between the LOD and the LOQ were assigned the mean value of LOD + LOQ (Figure S 3).

Samples with very low average DNA concentrations usually have an unproportionally high frequency of non-detects across technical replicates (McCall et al. 2014, Lesperance et al. 2021). To accurately estimate the average DNA concentration per eDNA sample it is important to assign a value of zero to true negatives, i.e., non-detects. We visually determined the average DNA concentration per *Bay* and *Visit* where the proportion of non-detects clearly deviated from the mean. This threshold was approximately at 8 copies μl^-1^ (Figure S 4). Hence non-detects below this threshold value were assigned a DNA concentration value of zero while values above the threshold, but below the LOD, were assigned a value of one-half of the LOD (Cohen and Ryan 1989).

### 2.3 Collection of angling data

In total, 24 coastal bays were fished at two occasions (visits), 8–20 days apart. Two of these bays were not sampled for eDNA due to logistic reasons (Villinge N 59° 5.7789’, E 18° 36.7948’, Jungfruskär, N 59° 8.4618’, E 18° 40.9969’). During each visit, the fishing was divided in two consecutive half days à 4 hours of active fishing each, alternating morning (08:00–12:00) and afternoon (13:00–17:00). In some cases, however, the fishing was instead performed during one full day (8 hours, Figure 1) due to logistics and weather conditions. The fishing was performed by six teams, each team consisted of two highly experienced pike anglers. The aim was to fish efficiently and catch as much fish as possible by choosing what the anglers considered to be the most suitable angling gear and bait. Sampling effort was quantified as rod hours, i.e., time fished per person. For each visit, the fishing teams recorded surface water temperature, number of seals at the site (estimated by eye), number of cormorants at the site (estimated by eye), numbers of other anglers present at the site (i.e. a boat with three anglers should be counted as three), and stationary fishing gear at the site (as indicated by buoys). Each pike caught was measured for total length using a tape measure, weight with a digital balance and sexed based on external characteristics (Casselman 1974). Spawning status was visually assessed according to expert judgment and classified as either pre-spawning (large girth indicating developed gonads but no running roe or milt), spawning (running roe or milt), post-spawning (spent fish, no running roe or milt and flaccid abdomen) or undefined (usually small fish without external characteristics indicating sexual maturation).

### 2.4 Abiotic data collection

Abiotic data was collected on each eDNA sampling visit to the bays using a Rinko ASTD-102 profiler (JFE Advantech Co., Ltd.). Depth, temperature, salinity, dissolved oxygen, turbidity and Chl-A-levels were measured from the surface to the bottom at the beginning and the end of each transect. The median value was calculated per depth profile and across transects to provide a grand median per bay visit.

### 2.5 Statistical analyses

Statistical analyses were performed using R, version 4.2.0 (R Core Team 2022) and the *tidyverse* suite of packages (Wickham et al. 2019). Linear mixed models were run using the *lme4* (Bates et al. 2015) and generalized linear models using the *MASS* package (Venables and Ripley 2002). Model evaluations were performed using the *DHARMa* (Hartig 2022) and *visreg* packages (Breheny and Burchett 2017) in combination with visual inspection of the residuals, outliers and leverage. Model fit was assessed using AICc (AIC corrected for sample size) and R square values were calculated with the *rsq* package (Zhang 2022). The level for statistical significance was set to α = 0.05. R-scripts and data for the analyses are provided as supplementary material.

#### 2.5.1 Standardization of angling data

To account for the potential influence of variables that might have affected catchability (i.e. rod-fishing efficiency) we ran a series of generalized linear mixed effects models to standardize the catch of pike in each bay and visit (hereafter called the CPUE-model). We modeled the number of pike caught per bay and visit using a poisson distribution and the log of fishing effort as an offset (n = 48). We used *Bay* and *Visit* nested within *Bay* as random factors on the intercept. The latter also functioned as an observation level random effect (OLRE) to handle overdispersion in the count data (Harrison 2014). For models that did not converge, the random effects were simplified to only include *Visit* nested within *Bay*. The number of other anglers (mean 3.0, range 0–25), number of cormorants (mean 9, range 0– 100), and water temperature (mean 7.9 °C, range 4–16) observed during angling were treated as continuous fixed effect variables. The other variables (number of seals and numbers of stationary fishing gear) were not included in the models since the data was too sporadic to be useful. Model selection consisted of fitting *i*) a base model with only random effects, *ii*) models with each fixed effect separately, *iii*) models with pairwise combinations of the fixed effects and *iv*) a full model with all variables, resulting in a total of 8 models. If two models were identified as equally parsimonious based on AICc, we chose the model with the strongest statistical significance for the fixed effects.

#### 2.5.2 Estimating spatiotemporal variation in eDNA concentrations

Because the eDNA and angling datasets were collected at different spatial scales, we modeled the average DNA concentration in each combination of bay and visit to make the two datasets compatible (hereafter called the eDNA-model). We used a generalized linear mixed model with a poisson distribution (Chambert et al. 2018). As response variable we used DNA copy number μl^-1^, the interaction between *Bay* and *Visit* was used as the fixed effect. Random effects on the intercept were transects nested within visits nested within bays and sample filter ID, which acted as an OLRE to account for the overdispersion in the data. We chose a mixed model over a conventional generalized linear model due to the hierarchical nesting of our data.

To assess the relative variance associated with either spatial or temporal variation we calculated the intra-class correlation coefficients (ICCs, or variance components) and their uncertainty (Nakagawa and Schielzeth 2010, Nakagawa et al. 2017). The model contained only random intercepts on *Bay, Bay: Visit, Bay: Visit: Transect, Bay: Visit: Transect:Filter* and the OLRE which consisted of each individual technical replicate in the qPCR dataset. Moreover, we fitted a second model, excluding the random effects for *Bay* and *Bay: Visit* and replaced them with the pooled effect of *Bay* and *Visit*. This was done in an attempt to account for the large uncertainty stemming from low within-level replication, especially at the finer scale such as within transects (two filters per transect).

#### 2.5.3 Modeling eDNA concentrations

To explain the variation in eDNA concentrations across bays and visits we tested and evaluated a range of generalized linear models. The response variable in these models was the average eDNA-concentration (DNA copies μl^-1^) estimated from the eDNA-model, which was assumed to come from a poisson distribution. Since the DNA concentrations were continuous but the poisson model requires integer values, we rounded the data up to the nearest integer (Chambert et al. 2018). As predictor variables we chose a range of variables known to affect the eDNA-signal. We chose *temperature* because it is a proximal variable that is intimately linked to physiological rate process in poikilotherms (Woods et al. 2003) and hence also DNA shedding and degradation (Jo et al. 2019), *bay size* because the eDNA concentration should be approximately proportional to the area/volume of a particular bay, assuming complete mixing of the water, i.e. a dilution effect (Yates et al. 2021b) and fish density estimated as CPUE from angling (Lacoursière-Roussel et al. 2016, Capo et al. 2019, Yates et al. 2021b, Stoeckle et al. 2021). Furthermore, because eDNA-concentrations have been shown to scale allometrically with fish size (Yates et al. 2021c, 2021a, Zhang et al. 2022) we also calculated the allometrically scaled average fish weight per bay and visit in the population as (ASM):

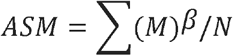

where *M* equals the individual weight (g), *β* equals a scaling coefficient (0.7) (Yates et al. 2022), and *N* the total number of fish caught per bay and visit. Effectively, this approach was a slight modification of the allometrically scaled mass (ASM) proposed by Yates et al. (2021b) and Stoeckle et al. (2021) since ASM in our case did not extend the calculation to the population level. Instead we used it as a covariate together with CPUE (sensu Spear et al. 2021). Another potential variable that could be expected to affect eDNA-concentrations is the spawning status of the population, since spawning events momentarily increase eDNA-levels due to increased activity of the fish (movements) but also release of gametes (Collins et al. 2022, Wu et al. 2022). This variable was strongly correlated with temperature and therefore omitted from the models by necessity. However, we calculated the variable *proportion spawned* to visualize the relationship between temperature and spawning status (Figure S 6),

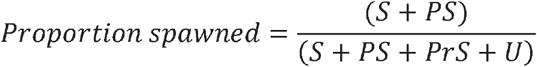

where *S* = spawning, *PS* = post-spawned, *PrS* = pre-spawning and *U* = undefined.

Based on the selected variables *temperature, bay size, CPUE* and *ASM* we used a forward selection process starting with the main effects of each variables, as well as the interaction between *CPUE* and *ASM* (Spear et al. 2021). After having selected the best fitting main effects, we included also potential interactions.

## 3 Results

### 3.1 qPCR data and quality control

Inhibition, operationally defined as a sample displaying Cq-values > 28.5 for the Cy5-labeled IPC was found in all or some of the technical replicates from five out of the 22 bays (“SP – Släpan/Ekefjärd”, “TT –Tomtviken/Urö”, “SS –Södersundet”, “MÖ –Möcklingeviken” and ÖL –Östra Lemaren”, Figure S 2). Consequently, these samples did not pass the quality control and were excluded from further analyses. In total, 285 filter samples (1135 samples including technical replicates) passed the quality control and were amenable for downstream analysis. Out of these samples (n = 1135), 61.9% were above the LOQ, 12.2% between LOD and LOQ, 2.2% below the LOD, and 23.6% were non-detects (Figure S 3). In order to not overestimate sample averages, 86.6% of the non-detects were imputed with zeros based on their overall high sample Cq-values (Figure S 4).

### 3.2 Descriptive abiotic data

The surveyed parts of the bays were shallow, with a median depth of 1 m (IQR = 0.7–1.4 m) across transect measurements. At the bay level, the temperature increased over the survey period from 3.2–11.7 °C (min-max). At the visit level the temperature increased from 4.9–8.3 °C on average. Salinity was relatively stable around 5 psu (median = 5.7, IQR = 5.1–5.9) but two bays situated in the inner most parts of the archipelago had lower salinity (SP – Släpan/Ekefjärd and MV –Myttingeviken, median = 2.4 and 2.6 psu respectively). These two bays were also characterized by markedly higher fluorescence intensities in the 640-980 nm range which is a proxy for Chlorophyll A concentrations (median = 13.8 and 5.9 ppb respectively relative the global median of 1.9 pbb).

### 3.3 Spatiotemporal eDNA dynamics

The amount of variance associated with the different levels of the eDNA survey could not be partitioned into clear spatial and temporal dynamics with the original model (Figure 3A). However, using the simplified model with the combined effect of *Bay:Visit*, differences emerged (Figure 3B). Surprisingly the variance within transects, i.e., between the two filters from the same collection of water, seemed to have a rather high variance (22 %, CI 14–33). The amount of variance explained at the local scale (within bays/visits, i.e., across transects) was lower (21 %, CI 11–34) than at the larger spatiotemporal scale between bays and visits (52 %, CI 33–64) (Figure 3B). This indicates that while significant, small scale spatial differences are of less importance compared to more large-scale and temporal processes during spring when pike spawn.

**Figure 3.**
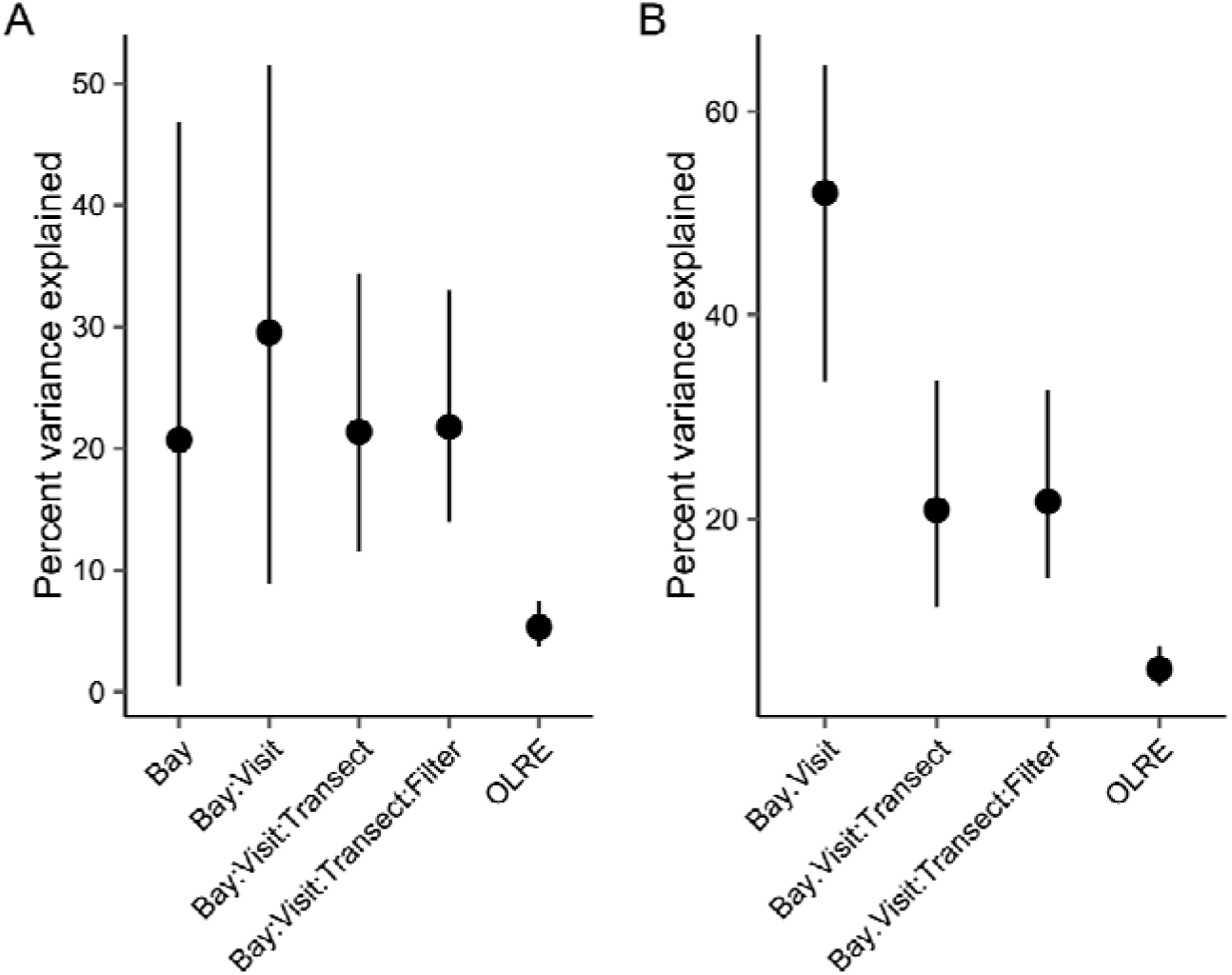
Intra class correlation coefficients for the random effects in the eDNA-model showing the percentage of variance explained in the model at different hierarchical levels from the full model (A) or a simplified version where the effects of Bay and Visit are pooled to one random effect (B).

### 3.4 Effects of fishing pressure on CPUE

The number of pike caught per rod-hour in the standardized angling was best explained by the negative effect of the number of other anglers present at the time of the survey (Table 1, model 4). It is worth noting that the negative effect of cormorants also appears important (Table 1). However, model 4 explained more of the fixed variance, had the lowest AICc and had a statistically significant predictor term (p = 0.023), which is why we chose this model to standardize the catches. The standardization model was used for the comparison with eDNA by predicting the catch (standardized pike abundance) in the absence of other anglers at an effort of 16 rod hours for each bay and visit.

**Table 1.**
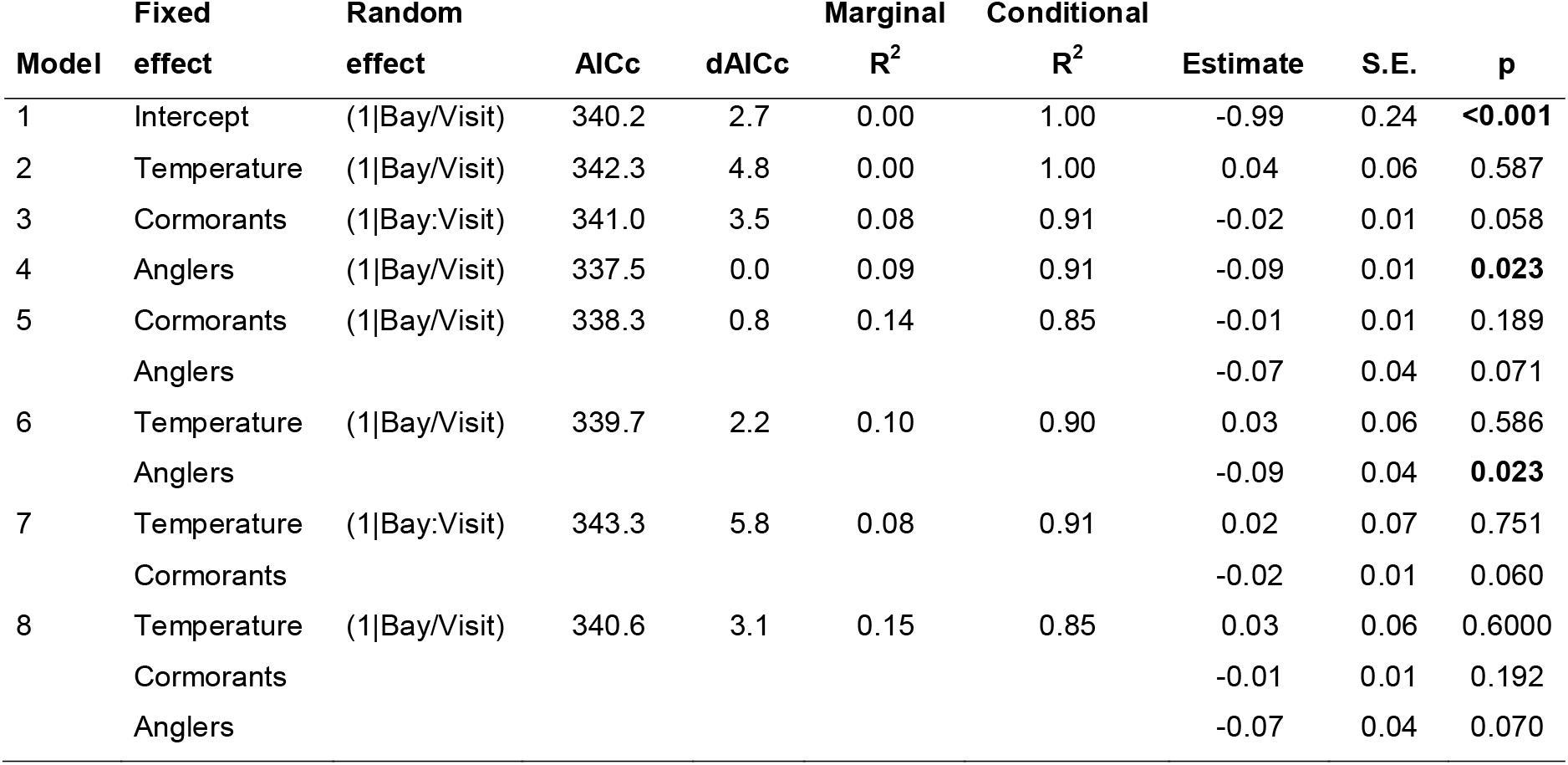
Model selection for the CPUE model. AICc is Akaike’s Information Criterion corrected for sample size and dAICc is the difference in AICc between a model and the best model. Marginal and conditional R^2^ show the proportion of variance explained by fixed factors only and total including random effects, respectively. Note that the conditional R^2^ is inflated by the use of observation level random effects. The estimate with associated standard error and p-value are given for each fixed effect. Significant p-values are highlighted in bold. Catch-Per-Unit-Effort (number of pikes caught per rod-hour) was used as the response variable in the models.

### 3.5 eDNA-abundance/biomass relationship

The forward selection process revealed that the eDNA concentration in the bays was primarily explained by temperature, followed by bay size (Table 2). Adding an interaction between temperature and bay size did not improve model fit, nor did adding either CPUE or ASM as main effects. However, the best fitting model included bay size and the interaction between temperature and CPUE (Model 18, Table 2, Table 3). Temperature and CPUE showed a significant and positive relationship with DNA concentration among bays and visits, which together with the negative effect of bay size explained 48% of the variance (Figure 4, Table 3).

**Figure 4.**
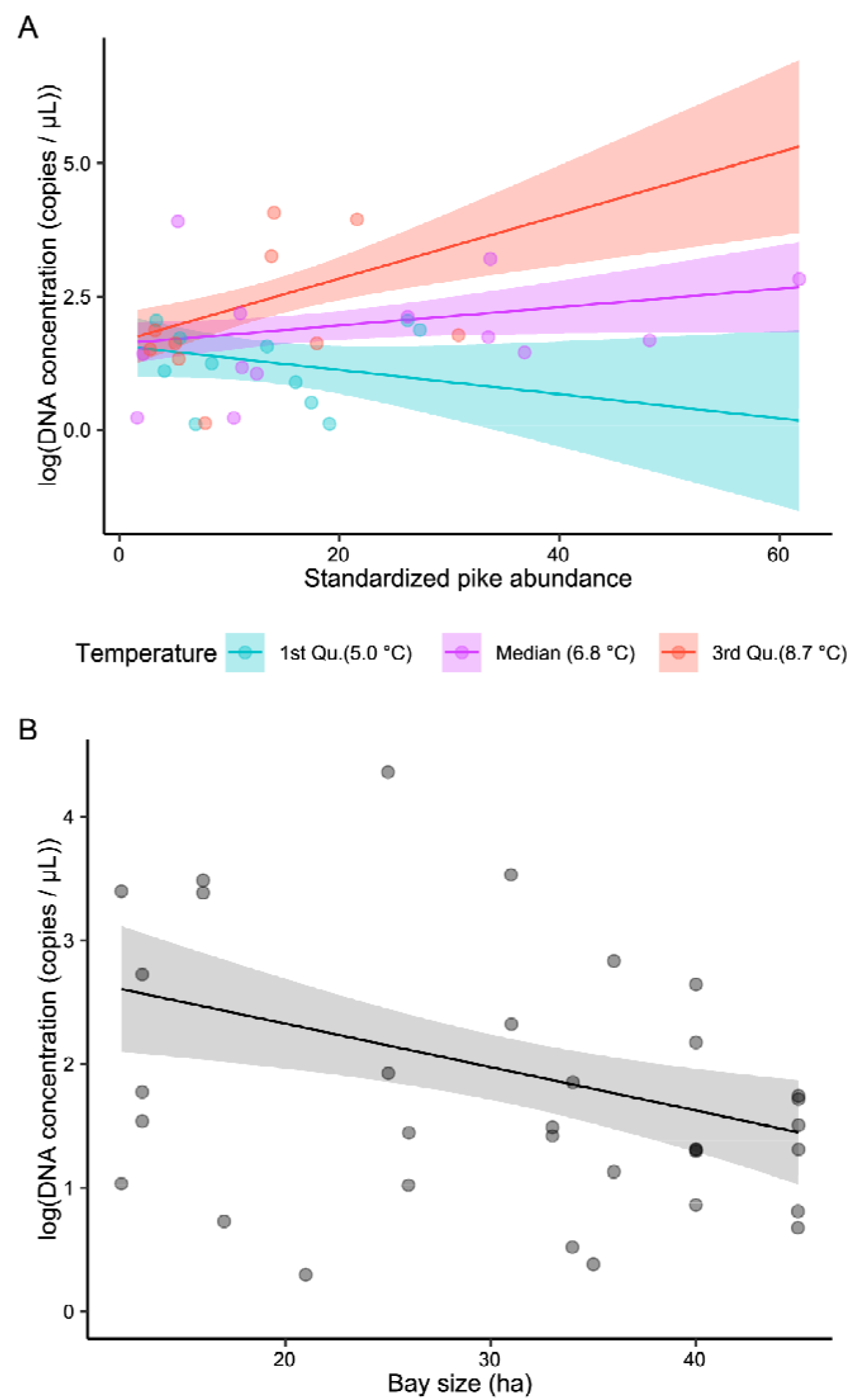
Model estimated DNA concentration (log-scale copies μL-1) as a function of A) standardized pike abundance, at different temperatures, and B) bay size (hectares). Points are partial residuals.R^2^ for the model was 0.48

**Table 2.**
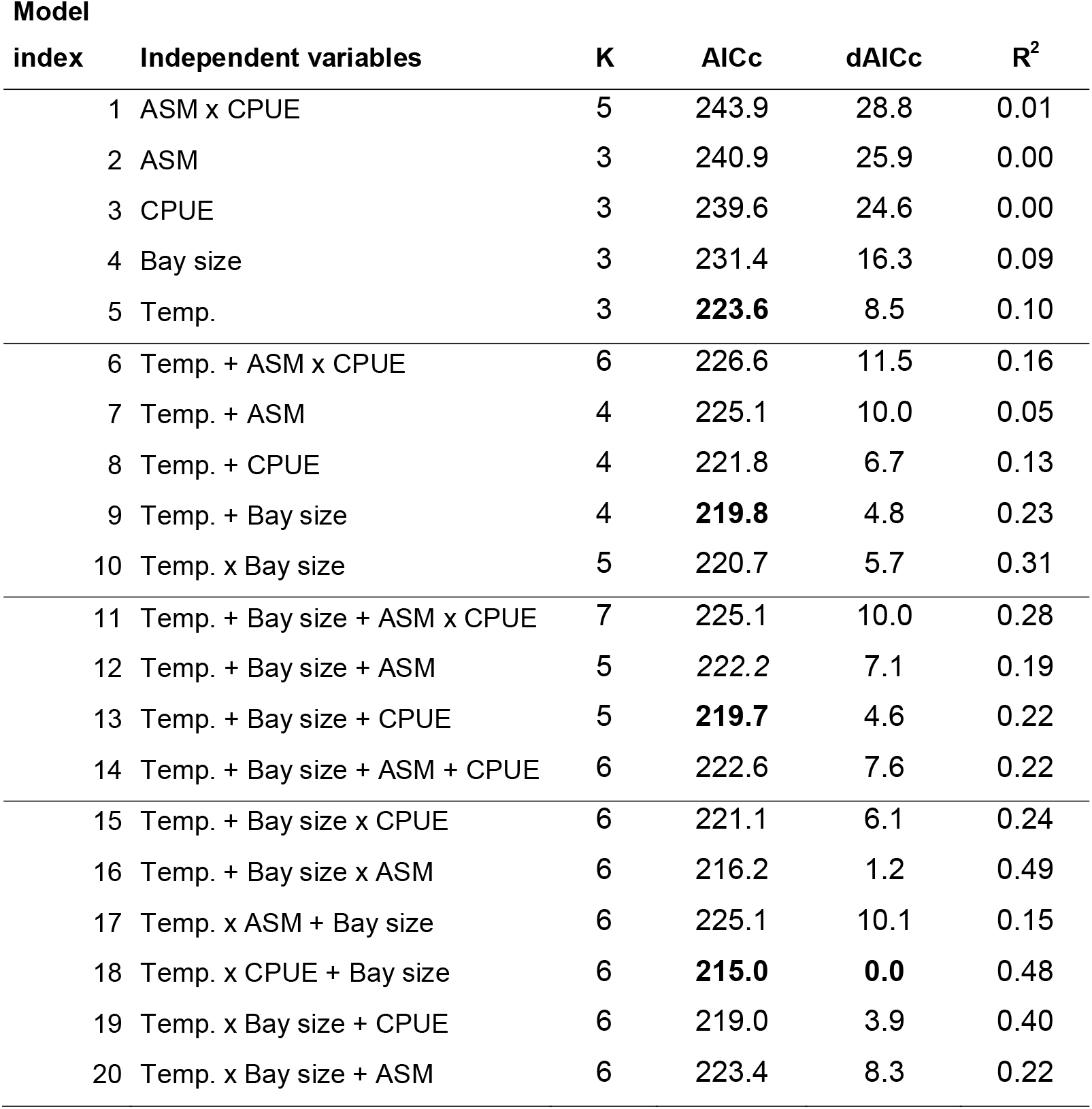
Selection of models explaining eDNA concentrations. The table is divided by a forward selection process. ASM is the allometrically scaled mean size in the population, CPUE is the standardized pike abundance, Temp. is water temperature and K is the number of parameters in the model. AICc is the Akaike Information Criterion corrected for sample size and dAICc is the difference in AICc between a model and the best model. R^2^ shows the proportion of variance explained by the model. Bold AICc indicates the best candidate model at each forward selection step.

**Table 3.** Model summary for the best performing model predicting eDNA concentrations (Model 18, Table 2). The model assumes a negative binomial distribution and uses a log-link function.

| Coefficients | Estimate | S.E | z | p-value | R<sup>2</sup> |
| --- | --- | --- | --- | --- | --- |
| Intercept | 2.65 | 0.98 | 2.702 | 0.007 | 0.48 |
| Temperature | 0.02 | 0.11 | 0.189 | 0.850 |  |
| CPUE | -0.13 | 0.05 | -2.446 | 0.014 |  |
| Bay size | -0.04 | 0.01 | -2.983 | 0.003 |  |
| Temperature × Bay size | 0.02 | 0.01 | 2.845 | 0.004 |  |

## 4 Discussion

As eDNA-abundance relationships are being established for many different species in a wide range of habitats, evidence is accumulating in favour of eDNA as a quantitative tool for monitoring fish populations. This suggests that the methodology bears potential for resource management and conservation purposes. However, the strength of these relationships has been variable, ranging from basically no relationship (Knudsen et al. 2019, Rourke et al. 2022) to rather high levels of correlation > 80% (Salter et al. 2019, Spear et al. 2021, Yates et al. 2021b). Reasons for this high level of heterogeneity are still not well understood but could be attributed to species-specific behavioural differences (Rourke et al. 2022), which makes it important to evaluate eDNA-abundance relationships at the species level (Jane et al. 2015, Lance and Guan 2020).

By comparing eDNA concentrations to standardized abundance metrics complemented by abiotic data we add to the growing line of evidence that eDNA can reflect the densities of wild fish populations and be a useful tool for monitoring fish populations. Specifically, our results show that eDNA can be applied to species that are generally undersampled by standard monitoring gear (gill-nets). Our results also show that this method can be used in semi-open habitats. However, the positive relationship between eDNA concentrations and the standardized pike abundance was not straightforward and we identified a number of confounding factors that will need to be taken in consideration as the eDNA methodology develops.

### 4.1. Spatiotemporal variation in eDNA concentrations

Using a high level of spatial replication within and across 22 bays we were able to assess the spatial variability of eDNA in a semi-enclosed coastal system. We found considerable variability across bay-visits but lower variation within bay-visits. Within specific bays, the variability in eDNA concentrations across transects was without typical patterns (Figure S 1, Figure S 5). We initially predicted that the central transect (D), which was situated in the deeper part of the bay, would consistently show lower DNA concentrations because it normally would fall outside the preferred vegetated habitat of spawning pike (Frost and Kipling 1967, Pursiainen et al. 2021). This was however not the case and we found no such discernible patterns.

The spatial distribution of eDNA has been shown to be patchy and vary seasonally in both marine and freshwater environments (Littlefair et al. 2021, Hervé et al. 2022). However, the degree to which concentrations vary mainly depends on the spatial distribution of the target species but also hydrographic and environmental conditions. For example, in larger lakes and marine systems it is common to find eDNA to be vertically stratified by thermoclines that form during periods of limited vertical mixing, effectively concentrating eDNA released from cold water species below the thermocline (Littlefair et al. 2021, Hervé et al. 2022). During our survey, the bays were thoroughly mixed which likely smoothed out any spatial differences (Table S 1). This lack of patchiness was also consistent over the two bay visits, albeit the average concentrations tended to be somewhat higher at the second visit as water temperatures rose (Figure S 1, Figure S 5, and Figure S 6). Moreover, our integrative approach of pooling water samples along the transects likely also contributed to decrease spatial patterns.

At smaller spatial scales, caging experiments have shown a rather limited detection distance in the range of 30 to 50 m in lakes (Dunker et al. 2016) and coastal waters (Murakami et al. 2019). In our case, we subsampled the transects with 50 m intervals and pooled the water at the end of each transect. In doing so we averaged out some level of variation making transects more similar to each other. Nevertheless, it should also be noted that the level of variation between filter replicates was of the same magnitude as across transects. This could partly be explained by low sample sizes and a statistical difficulty in partitioning the variance components but it could also be an effect of low DNA copy numbers and stochasticity which would warrant a higher level of in-field replication and filtration of larger volumes of water. Although the filtration of large water quantities may be cumbersome it may be performed using larger filter pore sizes. Such filters have a higher probability of capturing longer multi-copy nuclear eDNA fragments (Jo et al. 2020) which, due to their higher degradation rates compared to shorter mitochondrial DNA, better reflect instantaneous fish densities (Jo et al. 2022) and could possibly have improved the precision of our measurements.

### 4.2. eDNA-abundance relationship

#### 4.2.1. Temperature drives eDNA dynamics

Although we found a positive relationship between CPUE and eDNA concentrations we found an even stronger influence of temperature. Moreover, the effect of CPUE was only evident at higher temperatures suggesting either that *i*) pike abundance in the bays increased over the survey period and that there was an interaction with catchability (increased fish density but unchanged CPUE), *ii*) eDNA shedding rates increased with temperature (becoming detectable and fully quantifiable above a threshold temperature), and/or *iii*) that spawning, which increased with temperature (Figure S 6), had an additive effect.

We cannot rule out that pike abundance in the bays increased as the bays became warmer (i above) but temperature had no significant effect on CPUE (Table 1) suggesting the abundance of pike was relatively stable over the survey period. Moreover, catches were sometimes substantial already at temperatures as low as 3–4 °C (Figure 5) indicating that arrival to the spawning grounds happens well before temperatures have reached optimal spawning conditions which normally fall between 6–8 °C (Clark 1950, Frost and Kipling 1967). This is also supported by observations from other lake and river systems where arrival to the spawning grounds can precede the actual spawning event by several weeks or even months (Raat 1988). Therefore, direct effects of temperature on eDNA concentrations (ii and iii above) are more probable.

**Figure 5.**
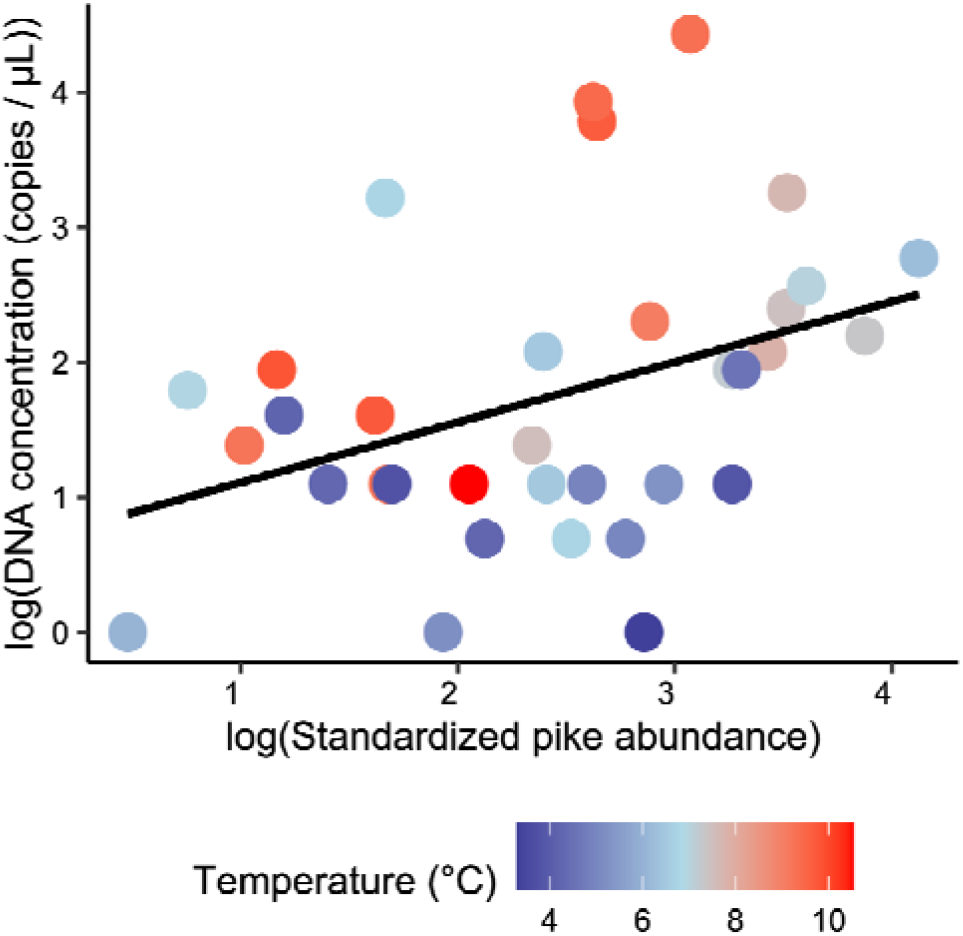
Raw data plot (log-log scale) of the relationship between DNA concentration and standardized pike abundance (CPUE) coloured by water temperature.

In line with our field observations, a laboratory study on brook trout has also shown a temperature mediated effect, resulting in a stronger eDNA-abundance/biomass relationship at higher temperatures (Lacoursière-Roussel et al. 2016). The authors suggested that the temperature-biomass interaction was driven by increased activity levels and metabolism. This is indeed very likely since metabolism and in extension DNA shedding rates are dependent on temperature (Kitchell et al. 1977, Bean 2010, Jo et al. 2019). Additional changes also take place as temperature rises, not the least an increase in the proportion of spawning fish (Figure S 6). As the fish spawn, their activity and physical interactions increase (Lucas 1992).

Simultaneously, the spawning event itself leads to the release of sperm which becomes readily incorporated in the eDNA pool (Tillotson et al. 2018, Tsuji and Shibata 2021, Holmes et al. 2022). The contribution of sperm could potentially be estimated by comparing ratios of nuclear and mitochondrial eDNA (Bylemans et al. (2017). Such comparisons rely on robust assays for both mitochondrial and nuclear DNA, but the latter is currently lacking for pike. In summary, we believe that temperature, especially during early spring in temperate regions, is a key driver affecting physiological processes such as metabolism and shedding rates, as well as behaviour and spawning activity–all of which has a strong influence on eDNA concentrations.

#### 4.2.2. CPUE based on angling likely underestimates true abundance

Even though we did our best to estimate the “true” pike abundance by modelling the effect of other anglers and deriving a standardized pike abundance, we found a relatively weak relationship between CPUE and eDNA concentrations. This could be a result of using angling data instead of census data from e.g., mark-recapture experiments (Spear et al. 2021). It is well known that angling success can vary due to local environmental conditions and, for species which are the target of catch-and-release practices like pike, also previous fishing intensity (Kuparinen et al. 2010, Arlinghaus et al. 2017, Chen and Zeng 2022). Even though we corrected for the number of anglers present during the rod-fishing, fishing pressure the days before remain unknown, which could also influence catchability. Similarly poor predictive capability of angler based abundance was obtained in a study investigating the eDNA-abundance relationships for brook charr (*Salvelinus fontinalis*, Mitchill 1814) in a series of Canadian lakes (Gaudet-Boulay et al. 2022). In that study, the CPUE of brook charr estimated from angling data predicted eDNA concentration in the lakes poorly, but the explanatory power of the model increased once the surface area of the lakes was accounted for, indicating that fish density measured per unit area is a better predictor (marginal R^2^ in models with only fish density as a predictor varied from 0.1–0.44). That observation is in accordance with our study where bay size as a covariate had a strong negative effect on eDNA concentrations (Table 2). Accounting for the size of the study area makes sense assuming that fish are heterogeneously distributed and concentrated to certain habitats. In the case of pike, it is very likely that most fish were aggregated close to the vegetated shore where spawning usually takes place (Clark 1950, Lucas 1992). Since the proportion of preferred habitat scales disproportionately with the square of bay area, and given that the eDNA is thoroughly mixed within the bay, this results in a dilution effect. Similar patterns of DNA dilution have been observed in rivers with elevated water discharges (Pont et al. 2022).

#### 4.2.3. Abundance or biomass as eDNA predictors?

Apart from using standardized abundance as a predictor of eDNA concentration we also tested to include allometrically scaled biomass (Yates et al. 2021c, 2022) by calculating the allometrically scaled mean population weight and using this as a covariate in our modelling. Several authors have recently shown improved relationships between fish biomass and eDNA concentrations when accounting for the size distribution of the fish community. Spear et al. (2021) saw an improvement in model *R^2^* from *0.62* to *0.81* when the mean size of walleye (*Sander vitreus*) was used as a covariate together with the estimated population biomass, while Yates et al. (2021b) saw an improvement in model R^2^ from *0.59* (fish/ha) and *0.63* (kg/ha) to *0.78* when accounting for allometric scaling in a study on brook charr. Although the evidence for allometric effects on eDNA production seem to be generalizable across species (Yates et al. 2022) we did not find a significant effect of including ASM in our pike models. The reason for this lack of effect is not clear but could potentially be attributed to a relatively homogeneous size distribution across bays. Indeed, the average pike weight per bay and visit in our study only differed by approximately a factor of three, while in the study by Yates et al. (2021b) the difference was substantially larger across lakes (factor ten difference). Furthermore, it is likely that the angling approach underestimated the abundance of smaller individuals which likely were not captured as efficiently by the anglers (x□ = 60 cm, SD = 10 cm). Such a size selectivity would effectively inflate the average size of the population, decrease the variance, and hence also influence allometry. Nevertheless, it is probable that allometric effects may be of greater importance during other seasons when local size distributions are more variable (Neumann and Willis 1995).

### 4.3. Conclusions

Our study supports the growing body of evidence showing a positive relationship between fish abundance/biomass and eDNA concentrations in the wild. Including abiotic data, we were able to explain nearly 50% of the variance in eDNA concentrations. This is in line with similar studies performed on other species and in different ecosystems (Yates et al. 2019). With the additional support from established eDNA-biomass relationships under more controlled conditions (Karlsson et al. 2022), it is likely that eDNA could be used to infer relative abundance data in wild pike populations. However, we also found temperature to be important, likely acting as a driver of fish activity and spawning that has a profound effect on eDNA concentrations. Temperatures that change rapidly, especially in temperate regions, will therefore induce unwanted variance, which may be difficult to account for. Hence, choosing appropriate sampling times will be crucial in order to make longitudinal data comparable. We therefore recommend that quantitative eDNA-surveys targeting northern pike should be performed at ambient temperatures > 10-12 °C when most pike have spawned but still have a high probability to stay aggregated in close proximity to their spawning areas.

## Supporting information

Supplementary maps

## Data archiving statement

The data and script for this study are available at Figshare, https://doi.org/10.6084/m9.figshare.21781622 following best practices (Roche et al. 2015), and was made available to editors and reviewers upon initial submission.

## Author contributions

All authors conceived and designed the study, EK, MO, GS, JS and PB collected the data, MO and GS conducted the statistical analyses and created the figures, MO and GS drafted the manuscript, with contributions from all co-authors. All authors approved the final submission.

## Ethics statement

All applicable international, national, and/or institutional guidelines for the care and use of animals were followed. The fish handled during the rod-fishing in this study complied with the standards and procedures stipulated by the Swedish Ministry of Agriculture and the ethical permit was approved by the Stockholm ethical committee (DNR S 33-15).

## Acknowledgements

We would like to thank Henrik C. Andersson at the Stockholm County Administrative board and all the anglers involved in the REFISK-project 2020 for providing the angling data. Dr Zandra Gerdes (Aquabiota Water Research), Ofir Svensson (Calluna AB) for help during laboratory work, and Ola Renman and John Persson (SLU) for help during the eDNA-data collection. This work was funded by a grant from the Swedish Environmental Protection Agency, Naturvårdsverket (NV-03728-17).

## References

Arlinghaus, R., J. Alós, T. Pieterek, and T. Klefoth. 2017. Determinants of angling catch of northern pike (Esox lucius) as revealed by a controlled whole-lake catch-and-release angling experiment—The role of abiotic and biotic factors, spatial encounters and lure type. Fisheries Research 186:648–657.

Arlinghaus, R., T. Klefoth, A. Kobler, and S. J. Cooke. 2008. Size Selectivity, Injury, Handling Time, and Determinants of Initial Hooking Mortality in Recreational Angling for Northern Pike: The Influence of Type and Size of Bait. North American Journal of Fisheries Management 28:123–134.

Bates, D., M. Mächler, B. Bolker, and S. Walker. 2015. Fitting Linear Mixed-Effects Models Using lme4. Journal of Statistical Software 67:1–48.

Bean, N. J. 2010. An Improved Bioenergetics Model for Northern Pike (Esox Lucius) of Box Canyon Reservoir, Pend Oreille River, Washington. Eastern Washington University.

Bergström, U., S. Larsson, M. Erlandsson, M. Ovegård, H. Ragnarsson Stabo, Ö. Östman, and G. Sundblad. 2022. Long-term decline in northern pike (Esox lucius L.) populations in the Baltic Sea revealed by recreational angling data. Fisheries Research 251:106307.

Bockrath, K. D., M. Tuttle-Lau, E. L. Mize, K. V. Ruden, and Z. Woiak. 2022. Direct comparison of eDNA capture and extraction methods through measuring recovery of synthetic DNA cloned into living cells. Environmental DNA 4:1000–1010.

Breheny, P., and W. Burchett. 2017. Visualization of Regression Models Using visreg. The R Journal 9:56–71.

Bylemans, J., E. M. Furlan, C. M. Hardy, P. McGuffie, M. Lintermans, and D. M. Gleeson. 2017. An environmental DNA-based method for monitoring spawning activity: a case study, using the endangered Macquarie perch (*Macquaria australasica*). Methods in Ecology and Evolution 8:646–655.

Capo, E., G. Spong, H. Königsson, and P. Byström. 2020. Effects of filtration methods and water volume on the quantification of brown trout (Salmo trutta) and Arctic char (Salvelinus alpinus) eDNA concentrations via droplet digital PCR. Environmental DNA 2:152–160.

Capo, E., G. Spong, S. Norman, H. Königsson, P. Bartels, and P. Byström. 2019. Droplet digital PCR assays for the quantification of brown trout (Salmo trutta) and Arctic char (Salvelinus alpinus) from environmental DNA collected in the water of mountain lakes. PLOS ONE 14:e0226638.

Casselman, J. M. 1974. External Sex Determination of Northern Pike, Esox lucius Linnaeus. Transactions of the American Fisheries Society 103:343–347.

Chambert, T., D. S. Pilliod, C. S. Goldberg, H. Doi, and T. Takahara. 2018. An analytical framework for estimating aquatic species density from environmental DNA. Ecology and Evolution 8:3468–3477.

Chen, L.-X., and L.-Q. Zeng. 2022. Previous experience alters individual vulnerability to angling of crucian carp (Carassius auratus). Behavioural Processes 195:104565.

Clark, C. F. 1950. Observations on the Spawning Habits of the Northern Pike, Esox lucius, in Northwestern Ohio. Copeia 1950:285.

Cohen, M. A., and P. B. Ryan. 1989. Observations Less than the Analytical Limit of Detection: A New Approach. JAPCA 39:328–329.

Collins, R. A., C. Baillie, N. C. Halliday, S. Rainbird, D. W. Sims, S. Mariani, and M. J. Genner. 2022. Reproduction influences seasonal eDNA variation in a temperate marine fish community. Limnology and Oceanography Letters 7:443–449.

Craig, J. F. 2008. A short review of pike ecology. Hydrobiologia 601:5–16.

Dejean, T., A. Valentini, A. Duparc, S. Pellier-Cuit, F. Pompanon, P. Taberlet, and C. Miaud. 2011. Persistence of Environmental DNA in Freshwater Ecosystems. PLOS ONE 6:e23398.

Diaz-Suarez, A., K. Noreikiene, V. Kisand, O. Burimski, R. Svirgsden, M. Rohtla, M. Ozerov, R. Gross, M. Vetemaa, and A. Vasemägi. 2022. Temporally stable small-scale genetic structure of Northern pike (Esox lucius) in the coastal Baltic Sea. Fisheries Research 254:106402.

Donadi, S., L. Bergström, J. M. Bertil Berglund, B. Anette, R. Mikkola, A. Saarinen, and U. Bergström. 2020. Perch and pike recruitment in coastal bays limited by stickleback predation and environmental forcing. Estuarine, Coastal and Shelf Science 246:107052.

Dunker, K. J., A. J. Sepulveda, R. L. Massengill, J. B. Olsen, O. L. Russ, J. K. Wenburg, and A. Antonovich. 2016. Potential of Environmental DNA to Evaluate Northern Pike (Esox lucius) Eradication Efforts: An Experimental Test and Case Study. PloS One 11:e0162277.

Eklöf, J. S., G. Sundblad, M. Erlandsson, S. Donadi, J. P. Hansen, B. K. Eriksson, and U. Bergström. 2020. A spatial regime shift from predator to prey dominance in a large coastal ecosystem. Communications Biology 3:1–9.

Forsman, A., P. Tibblin, H. Berggren, O. Nordahl, P. Koch-Schmidt, and P. Larsson. 2015. Pike *Esox lucius* as an emerging model organism for studies in ecology and evolutionary biology: a review: *esox lucius* as a model in ecology and evolution. Journal of Fish Biology 87:472–479.

Frost, W. E., and C. Kipling. 1967. A Study of Reproduction, Early Life, Weight-Length Relationship and Growth of Pike, Esox lucius L., in Windermere. The Journal of Animal Ecology 36:651.

Gaudet-Boulay, M., E. García-Machado, M. Laporte, M. Yates, B. Bougas, C. Hernandez, G. Côté, A. Gilbert, and L. Bernatchez. 2022. Relationship between brook charr (Salvelinus fontinalis) eDNA concentration and angling data in structured wildlife areas. Environmental DNA n/a.

Hansson, S., U. Bergström, E. Bonsdorff, T. Härkönen, N. Jepsen, L. Kautsky, K. Lundström, S.-G. Lunneryd, M. Ovegård, J. Salmi, D. Sendek, and M. Vetemaa. 2017. Competition for the fish – fish extraction from the Baltic Sea by humans, aquatic mammals, and birds. ICES Journal of Marine Science.

Hartig, F. 2022. DHARMa: Residual Diagnostics for Hierarchical (Multi-Level / Mixed) Regression Models.

Hernandez, C., B. Bougas, A. Perreault-Payette, A. Simard, G. Côté, and L. Bernatchez. 2020. 60 specific eDNA qPCR assays to detect invasive, threatened, and exploited freshwater vertebrates and invertebrates in Eastern Canada. Environmental DNA 2:373–386.

Hervé, A., I. Domaizon, J.-M. Baudoin, T. Dejean, P. Gibert, P. Jean, T. Peroux, J.-C. Raymond, A. Valentini, M. Vautier, and M. Logez. 2022. Spatio-temporal variability of eDNA signal and its implication for fish monitoring in lakes. PLOS ONE 17:e0272660.

Holmes, V., J. Aman, G. York, and M. T. Kinnison. 2022. Environmental DNA detects Spawning Habitat of an ephemeral migrant fish (Anadromous Rainbow Smelt: Osmerus mordax). BMC Ecology and Evolution 22:121.

Jane, S. F., T. M. Wilcox, K. S. McKelvey, M. K. Young, M. K. Schwartz, W. H. Lowe, B. H. Letcher, and A. R. Whiteley. 2015. Distance, flow and PCR inhibition: eDNA dynamics in two headwater streams. Molecular Ecology Resources 15:216–227.

Jo, T., H. Murakami, R. Masuda, and T. Minamoto. 2020. Selective collection of long fragments of environmental DNA using larger pore size filter. Science of The Total Environment 735:139462.

Jo, T., H. Murakami, S. Yamamoto, R. Masuda, and T. Minamoto. 2019. Effect of water temperature and fish biomass on environmental DNA shedding, degradation, and size distribution. Ecology and Evolution 9:1135–1146.

Jo, T., K. Takao, and T. Minamoto. 2022. Linking the state of environmental DNA to its application for biomonitoring and stock assessment: Targeting mitochondrial/nuclear genes, and different DNA fragment lengths and particle sizes. Environmental DNA 4:271–283.

Kačergytė, I., E. Petersson, D. Arlt, M. Hellström, J. Knape, J. Spens, M. Żmihorski, and T. Pärt. 2021. Environmental DNA metabarcoding elucidates patterns of fish colonisation and co-occurrences with amphibians in temperate wetlands created for biodiversity. Freshwater Biology 66:1915–1929.

Karlsson, E., M. Ogonowski, G. Sundblad, J. Sundin, O. Svensson, I. Nousiainen, and A. Vasemägi. 2022. Strong positive relationships between eDNA concentrations and biomass in juvenile and adult pike (Esox lucius) under controlled conditions: Implications for monitoring. Environmental DNA 4:881–893.

Kitchell, J. F., D. J. Stewart, and D. Weininger. 1977. Applications of a Bioenergetics Model to Yellow Perch (Perca flavescens) and Walleye (Stizostedion vitreum vitreum). Journal of the Fisheries Research Board of Canada 34:1922–1935.

Klymus, K. E., C. M. Merkes, M. J. Allison, C. S. Goldberg, C. C. Helbing, M. E. Hunter, C. A. Jackson, R. F. Lance, A. M. Mangan, E. M. Monroe, A. J. Piaggio, J. P. Stokdyk, C. C. Wilson, and C. A. Richter. 2020. Reporting the limits of detection and quantification for environmental DNA assays. Environmental DNA 2:271–282.

Knudsen, S. W., R. B. Ebert, M. Hesselsøe, F. Kuntke, J. Hassingboe, P. B. Mortensen, P. F. Thomsen, E. E. Sigsgaard, B. K. Hansen, E. E. Nielsen, and P. R. Møller. 2019. Species-specific detection and quantification of environmental DNA from marine fishes in the Baltic Sea. Journal of Experimental Marine Biology and Ecology 510:31–45.

Kuparinen, A., T. Klefoth, and R. Arlinghaus. 2010. Abiotic and fishing-related correlates of angling catch rates in pike (Esox lucius). Fisheries Research 105:111–117.

Lacoursière-Roussel, A., M. Rosabal, and L. Bernatchez. 2016. Estimating fish abundance and biomass from eDNA concentrations: variability among capture methods and environmental conditions. Molecular Ecology Resources 16:1401–1414.

Laikre, L., L. M. Miller, A. Palmé, S. Palm, A. R. Kapuscinski, G. Thoresson, and N. Ryman. 2005. Spatial genetic structure of northern pike (Esox lucius) in the Baltic Sea. Molecular Ecology 14:1955–1964.

Lance, R. F., and X. Guan. 2020. Variation in inhibitor effects on qPCR assays and implications for eDNA surveys. Canadian Journal of Fisheries and Aquatic Sciences 77:23–33.

Lesperance, M. L., M. J. Allison, L. C. Bergman, M. D. Hocking, and C. C. Helbing. 2021. A statistical model for calibration and computation of detection and quantification limits for low copy number environmental DNA samples. Environmental DNA 3:970–981.

Li, C., H. Long, S. Yang, Y. Zhang, F. Tang, W. Jin, G. Wang, W. Chang, Y. Pi, L. Gao, L. Ma, M. Zhao, H. Zheng, Y. Gong, Y. Liu, and K. Jiang. 2022. eDNA assessment of pelagic fish diversity, distribution, and abundance in the central Pacific Ocean. Regional Studies in Marine Science 56:102661.

Littlefair, J. E., L. E. Hrenchuk, P. J. Blanchfield, M. D. Rennie, and M. E. Cristescu. 2021. Thermal stratification and fish thermal preference explain vertical eDNA distributions in lakes. Molecular Ecology 30:3083–3096.

Lucas, M. C. 1992. Spawning activity of male and female pike, Esox lucius L., determined by acoustic tracking. Canadian Journal of Zoology 70:191–196.

McCall, M. N., H. R. McMurray, H. Land, and A. Almudevar. 2014. On non-detects in qPCR data. Bioinformatics (Oxford, England) 30:2310–2316.

Möller, S., H. M. Winkler, S. Richter, and R. Bastrop. 2021. Genetic population structure of pike (Esox lucius Linnaeus, 1758) in the brackish lagoons of the southern Baltic Sea. Ecology of Freshwater Fish 30:140–149.

Murakami, H., S. Yoon, A. Kasai, T. Minamoto, S. Yamamoto, M. K. Sakata, T. Horiuchi, H. Sawada, M. Kondoh, Y. Yamashita, and R. Masuda. 2019. Dispersion and degradation of environmental DNA from caged fish in a marine environment. Fisheries Science 85:327–337.

Nakagawa, S., P. C. D. Johnson, and H. Schielzeth. 2017. The coefficient of determination R2 and intra-class correlation coefficient from generalized linear mixed-effects models revisited and expanded. Journal of The Royal Society Interface 14:20170213.

Nakagawa, S., and H. Schielzeth. 2010. Repeatability for Gaussian and non-Gaussian data: a practical guide for biologists. Biological Reviews 85:935–956.

Neumann, R. M., and D. W. Willis. 1995. Seasonal Variation in Gill-Net Sample Indexes for Northern Pike Collected from a Glacial Prairie Lake. North American Journal of Fisheries Management 15:838–844.

Olsen, J. B., C. J. Lewis, R. L. Massengill, K. J. Dunker, and J. K. Wenburg. 2015. An evaluation of target specificity and sensitivity of three qPCR assays for detecting environmental DNA from Northern Pike (<Emphasis Type=“Italic”>Esox lucius</Emphasis>). Conservation Genetics Resources 7:615–617.

Olsen, J. B., C. J. Lewis, R. L. Massengill, K. J. Dunker, and J. K. Wenburg. 2016. Erratum to: An evaluation of target specificity and sensitivity of three qPCR assay for detecting environmental DNA from Northern Pike (Esox lucius). Conservation Genetics Resources 8:89–89.

Pont, D., P. Meulenbroek, V. Bammer, T. Dejean, T. Erős, P. Jean, M. Lenhardt, C. Nagel, L. Pekarik, M. Schabuss, B. C. Stoeckle, E. Stoica, H. Zornig, A. Weigand, and A. Valentini. 2022. Quantitative monitoring of diverse fish communities on a large scale combining eDNA metabarcoding and qPCR. Molecular Ecology Resources n/a:1–14.

Pursiainen, A., L. Veneranta, S. Kuningas, A. Saarinen, and M. Kallasvuo. 2021. The more sheltered, the better – Coastal bays and lagoons are important reproduction habitats for pike in the northern Baltic Sea. Estuarine, Coastal and Shelf Science 259:107477.

R Core Team. 2022. R: A Language and Environment for Statistical Computing. R Foundation for Statistical Computing, Vienna, Austria.

Raat, A. J. P. 1988. Synopsis of Biological Data on the Northern Pike Esox lucius Linnaeus, 1758. Page 178. Synopsis, FAO, Rome.

Roche, D. G., L. E. B. Kruuk, R. Lanfear, and S. A. Binning. 2015. Public Data Archiving in Ecology and Evolution: How Well Are We Doing? PLOS Biology 13:e1002295.

Rourke, M. L., A. M. Fowler, J. M. Hughes, M. K. Broadhurst, J. D. DiBattista, S. Fielder, J. Wilkes Walburn, and E. M. Furlan. 2021. Environmental DNA (eDNA) as a tool for assessing fish biomass: A review of approaches and future considerations for resource surveys. Environmental DNA n/a.

Rourke, M. L., J. W. Walburn, M. K. Broadhurst, A. M. Fowler, J. M. Hughes, D. S. Fielder, J. D. DiBattista, and E. M. Furlan. 2022. Poor utility of environmental DNA for estimating the biomass of a threatened freshwater teleost; but clear direction for future candidate assessments. Fisheries Research:106545.

Salter, I., M. Joensen, R. Kristiansen, P. Steingrund, and P. Vestergaard. 2019. Environmental DNA concentrations are correlated with regional biomass of Atlantic cod in oceanic waters. Communications Biology 2:1–9.

Shelton, A. O., A. Ramón-Laca, A. Wells, J. Clemons, D. Chu, B. E. Feist, R. P. Kelly, S. L. Parker-Stetter, R. Thomas, K. M. Nichols, and L. Park. 2022. Environmental DNA provides quantitative estimates of Pacific hake abundance and distribution in the open ocean. Proceedings of the Royal Society B: Biological Sciences 289:20212613.

Song, J. W., M. J. Small, and E. A. Casman. 2017. Making sense of the noise: The effect of hydrology on silver carp eDNA detection in the Chicago area waterway system. Science of The Total Environment 605–606:713–720.

Spear, M. J., H. S. Embke, P. J. Krysan, and M. J. V. Zanden. 2021. Application of eDNA as a tool for assessing fish population abundance. Environmental DNA 3:83–91.

Stoeckle, B. C., S. Beggel, A. F. Cerwenka, E. Motivans, R. Kuehn, and J. Geist. 2017. A systematic approach to evaluate the influence of environmental conditions on eDNA detection success in aquatic ecosystems. PloS One 12:e0189119.

Stoeckle, M. Y., J. Adolf, Z. Charlop-Powers, K. J. Dunton, G. Hinks, and S. M. VanMorter. 2021. Trawl and eDNA assessment of marine fish diversity, seasonality, and relative abundance in coastal New Jersey, USA. ICES Journal of Marine Science 78:293–304.

Sundblad, G., and U. Bergström. 2014. Shoreline development and degradation of coastal fish reproduction habitats. Ambio 43:1020–1028.

Svensson, R. 2021. Development of northern pike (Esox lucius) populations in the Baltic Sea, and potential effects of grey seal (Halichoerus grypus) predation. Master thesis in Biology, Swedish University of Agricultural Sciences, Öregrund, Sweden.

Takahara, T., T. Minamoto, and H. Doi. 2013. Using Environmental DNA to Estimate the Distribution of an Invasive Fish Species in Ponds. PLOS ONE 8:e56584.

Thomsen, P. F., J. Kielgast, L. L. Iversen, P. R. Møller, M. Rasmussen, and E. Willerslev. 2012. Detection of a Diverse Marine Fish Fauna Using Environmental DNA from Seawater Samples. PLOS ONE 7:e41732.

Tillotson, M. D., R. P. Kelly, J. J. Duda, M. Hoy, J. Kralj, and T. P. Quinn. 2018. Concentrations of environmental DNA (eDNA) reflect spawning salmon abundance at fine spatial and temporal scales. Biological Conservation 220:1–11.

Tsuji, S., and N. Shibata. 2021. Identifying spawning events in fish by observing a spike in environmental DNA concentration after spawning. Environmental DNA 3:190–199.

Venables, W. N., and B. D. Ripley. 2002. Modern Applied Statistics with S. Fourth. Springer, New York.

Villegas-Ríos, D., J. Alós, M. Palmer, S. Lowerre-Barbieri, R. Bañón, A. Alonso-Fernández, and F. Saborido-Rey. 2014. Life-history and activity shape catchability in a sedentary fish. Marine Ecology Progress Series 515:239–250.

Wennerström, L., J. Olsson, N. Ryman, and L. Laikre. 2016. Temporally stable, weak genetic structuring in brackish water northern pike (*Esox lucius*) in the Baltic Sea indicates a contrasting divergence pattern relative to freshwater populations. Canadian Journal of Fisheries and Aquatic Sciences:1–10.

Wickham, H., M. Averick, J. Bryan, W. Chang, L. D. McGowan, R. François, G. Grolemund, A. Hayes, L. Henry, J. Hester, M. Kuhn, T. L. Pedersen, E. Miller, S. M. Bache, K. Müller, J. Ooms, D. Robinson, D. P. Seidel, V. Spinu, K. Takahashi, D. Vaughan, C. Wilke, K. Woo, and H. Yutani. 2019. Welcome to the tidyverse. Journal of Open Source Software 4:1686.

Wilcox, T. M., K. S. McKelvey, M. K. Young, A. J. Sepulveda, B. B. Shepard, S. F. Jane, A. R. Whiteley, W. H. Lowe, and M. K. Schwartz. 2016. Understanding environmental DNA detection probabilities: A case study using a stream-dwelling char Salvelinus fontinalis. Biological Conservation 194:209–216.

Woods, H. A., W. Makino, J. B. Cotner, S. E. Hobbie, J. F. Harrison, K. Acharya, and J. J. Elser. 2003. Temperature and the chemical composition of poikilothermic organisms. Functional Ecology 17:237–245.

Wu, L., Q. Wu, T. Inagawa, J. Okitsu, S. Sakamoto, and T. Minamoto. 2022. Estimating the spawning activity of fish species using nuclear and mitochondrial environmental DNA concentrations and their ratios. Freshwater Biology n/a.

Yates, M. C., M. E. Cristescu, and A. M. Derry. 2021a. Integrating physiology and environmental dynamics to operationalize environmental DNA (eDNA) as a means to monitor freshwater macro-organism abundance. Molecular Ecology 30:6531–6550.

Yates, M. C., D. J. Fraser, and A. M. Derry. 2019. Meta-analysis supports further refinement of eDNA for monitoring aquatic species-specific abundance in nature. Environmental DNA 1:5–13.

Yates, M. C., D. M. Glaser, J. R. Post, M. E. Cristescu, D. J. Fraser, and A. M. Derry. 2021b. The relationship between eDNA particle concentration and organism abundance in nature is strengthened by allometric scaling. Molecular Ecology 30:3068–3082.

Yates, M. C., T. M. Wilcox, K. S. McKelvey, M. K. Young, M. K. Schwartz, and A. M. Derry. 2021c. Allometric scaling of eDNA production in stream-dwelling brook trout (Salvelinus fontinalis) inferred from population size structure. Environmental DNA 3:553–560.

Yates, M. C., T. W. Wilcox, M. Y. Stoeckle, and D. D. Heath. 2022, April 22. Interspecific allometric scaling in eDNA production in fishes reflects physiological and surface area allometry. bioRxiv.

Zhang, D. 2022. rsq: R-Squared and Related Measures.

Zhang, J., R. Ding, Y. Wang, and J. Wen. 2022. Experimental study on the response relationship between environmental DNA concentration and biomass of Schizothorax prenanti in still water. Frontiers in Ecology and Evolution 10.

