## Supporting information for "Temperature moderates eDNA-biomass relationships in northern pike"

**Figures**

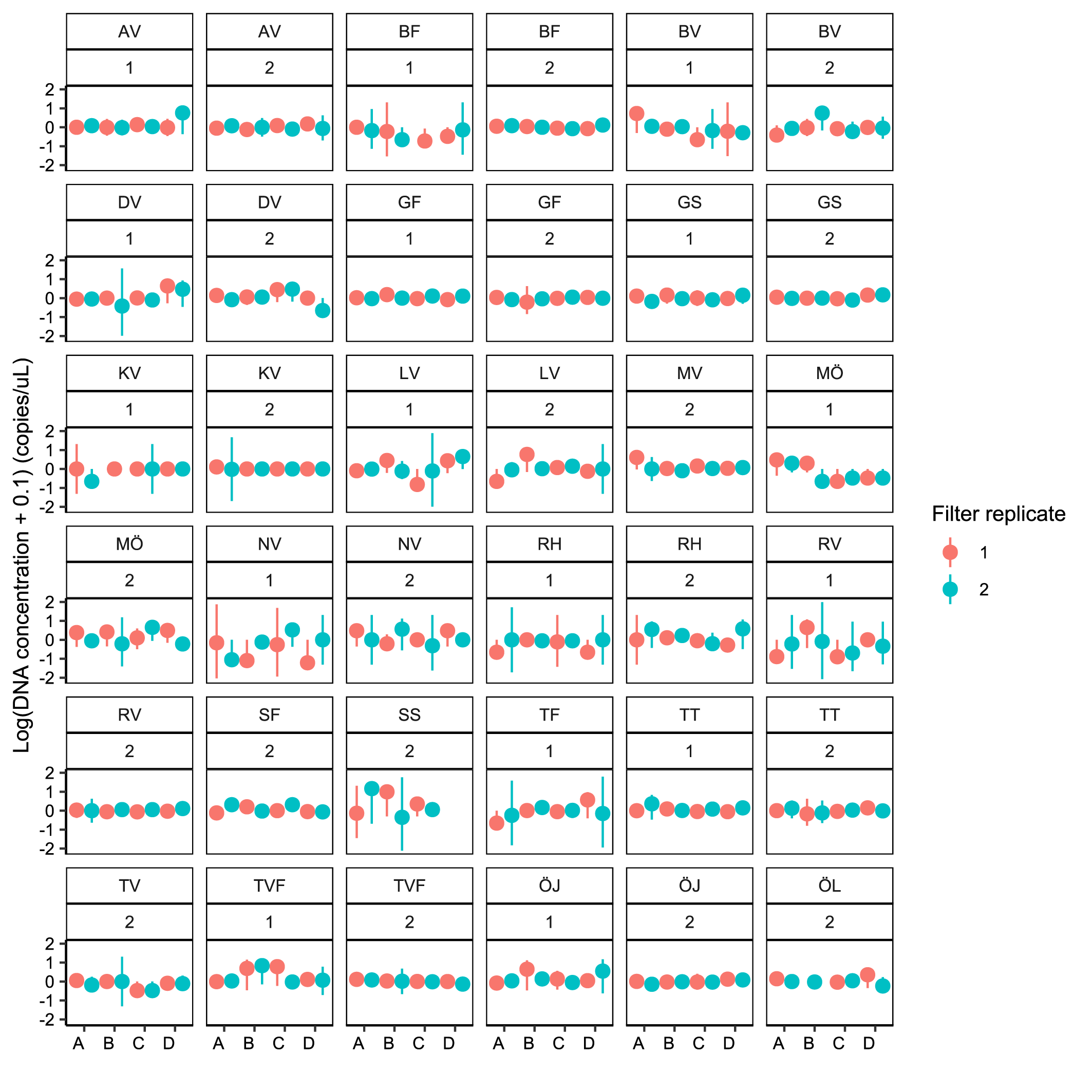

*Figure S 1. Filter plot showing Log transformed eDNA concentrations (copies µL^-1^) per bay, visit (1 and 2), transect (A-D) and filter replicate (1 and 2). Each filter consists of four technical qPCR replicates. Bay ID: AV = Askviken, BF = Björnöfjärden/Torpe Infjärd, BV = Byviken/Ryssundet, DV = Dalviken, GF = Gisslingöfladen, GS = Granösundet, KV = Kyrkviken/Utö, LV = Lännåkersviken, MV = Myttingeviken, MÖ = Mulö/Lögla, NV = Nynäsviken, RH = Rotholmaviken, RV = Rassa vikar, SF = Söderöfjärden/Sladdarön, SP = Släpan/Ekefjärd, SS = Södersundet, TF = Tofladen/Gropaviken, TT = Tomtviken/Urö, TV = Tranvik/Djuröviken, TVF = Tranviksfjärden, ÖJ = Öjaren/Söderöra/Norröra, ÖL = Östra Lemaren.*

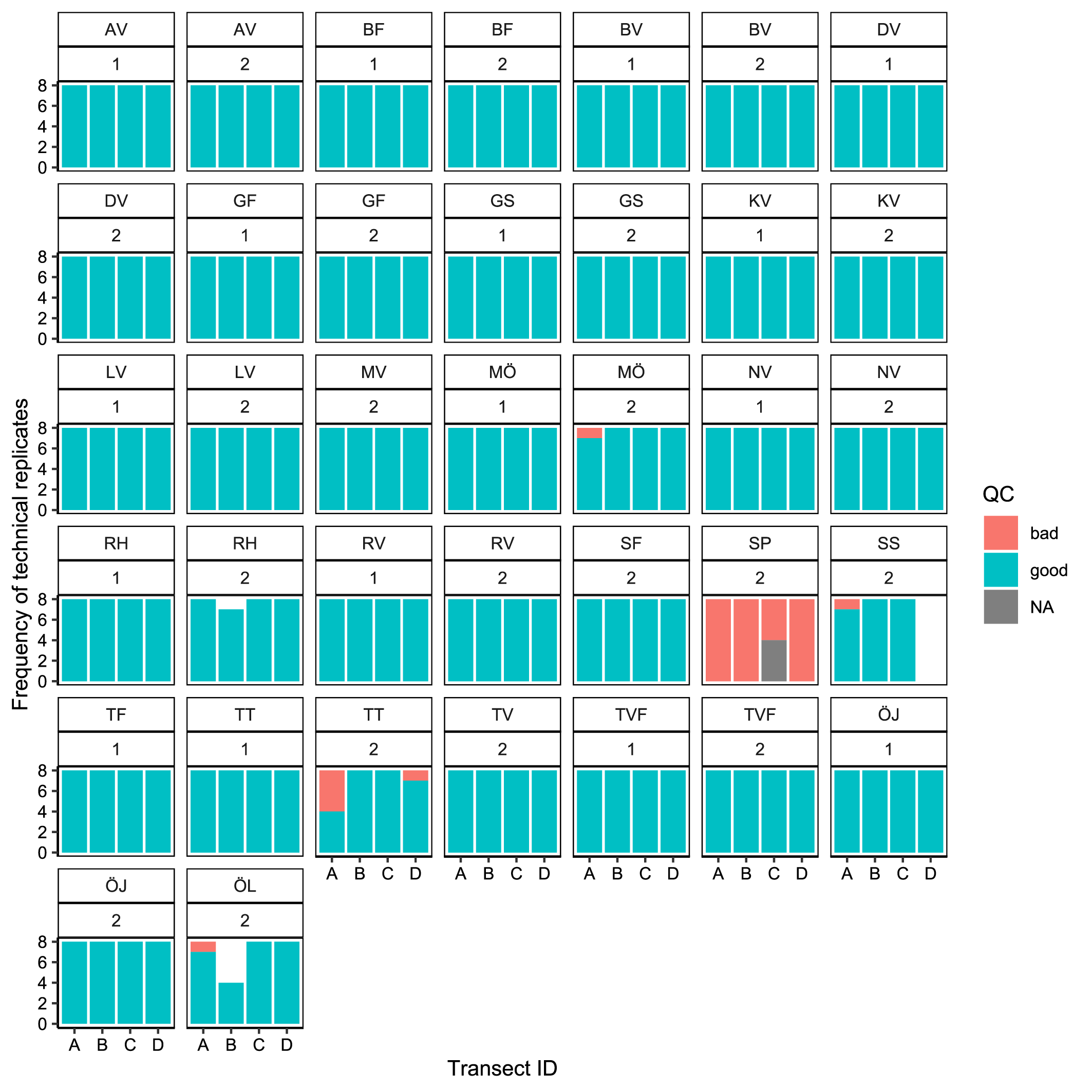

*Figure S 2. Barplot showing the frequency of technical qPCR replicates flagged for inhibition by having Cq values > 28.5 for the internal positive control. Data is divided per bay, visit and transect. Bay ID: AV = Askviken, BF = Björnöfjärden/Torpe Infjärd, BV = Byviken/Ryssundet, DV = Dalviken, GF = Gisslingöfladen, GS = Granösundet, KV = Kyrkviken/Utö, LV = Lännåkersviken, MV = Myttingeviken, MÖ = Mulö/Lögla, NV = Nynäsviken, RH = Rotholmaviken, RV = Rassa vikar, SF = Söderöfjärden/Sladdarön, SP = Släpan/Ekefjärd, SS = Södersundet, TF = Tofladen/Gropaviken, TT = Tomtviken/Urö, TV = Tranvik/Djuröviken, TVF = Tranviksfjärden, ÖJ = Öjaren/Söderöra/Norröra, ÖL = Östra Lemaren*

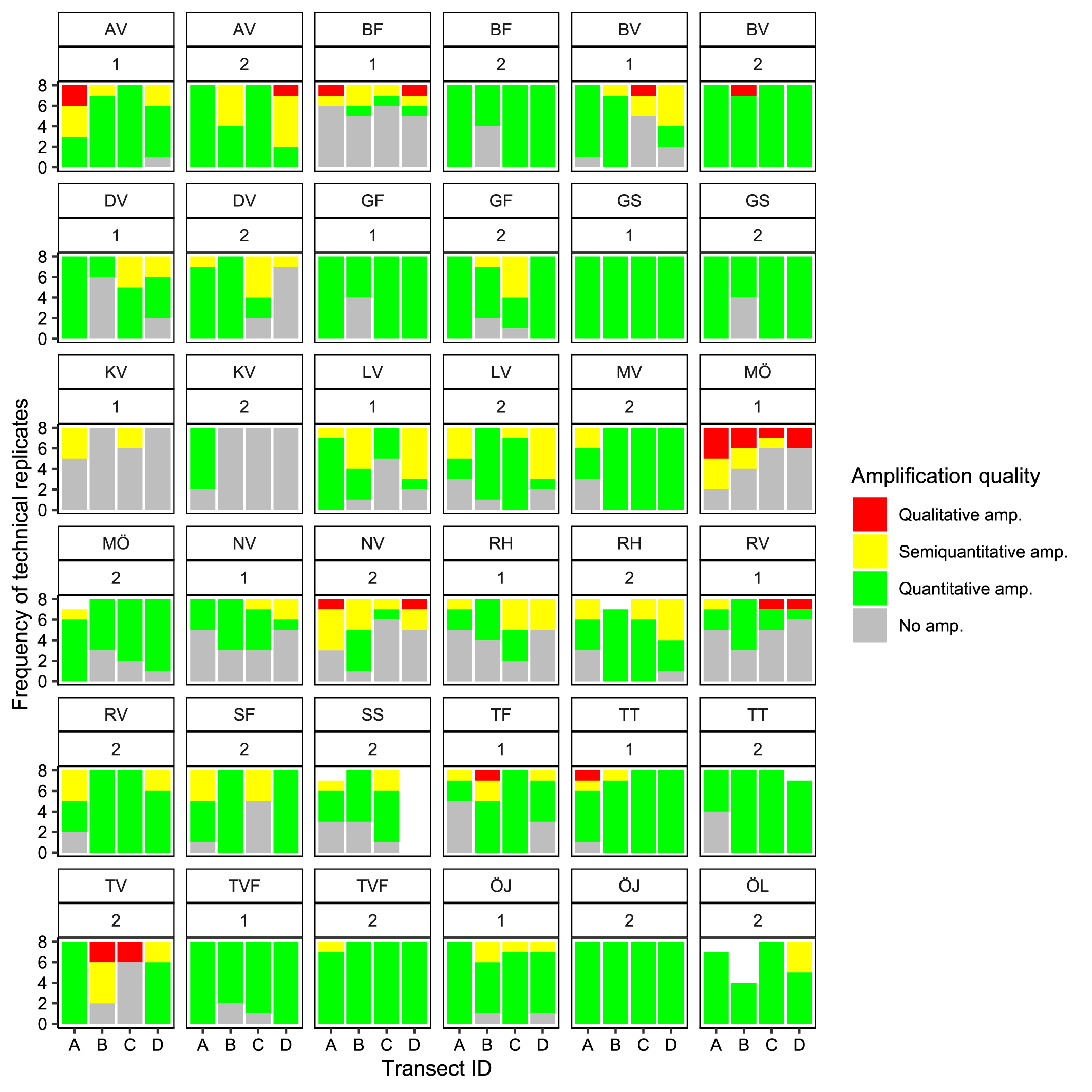

Figure S 3. Barplot showing the frequency of technical qPCR replicates classified as either quantitative (red bars, values above the limit of quantification –LOQ), semi-quantitative (yellow bars, between LOQ and LOD –the limit of detection), qualitative (green bars, above LOD but below LOQ), and non-detects (grey bars, No. amp). Data is divided per bay, visit and transect. Bay ID: AV = Askviken, BF = Björnöfjärden/Torpe Infjärd, BV = Byviken/Ryssundet, DV = Dalviken, GF = Gisslingöfladen, GS = Granösundet, KV = Kyrkviken/Utö, LV = Lännåkersviken, MV = Myttingeviken, MÖ = Mulö/Lögla, NV = Nynäsviken, RH = Rotholmaviken, RV = Rassa vikar, SF = Söderöfjärden/Sladdarön, SP = Släpan/Ekefjärd, SS = Södersundet, TF = Tofladen/Gropaviken, TT = Tomtviken/Urö, TV = Tranvik/Djuröviken, TVF = Tranviksfjärden, ÖJ = Öjaren/Söderöra/Norröra, ÖL = Östra Lemaren

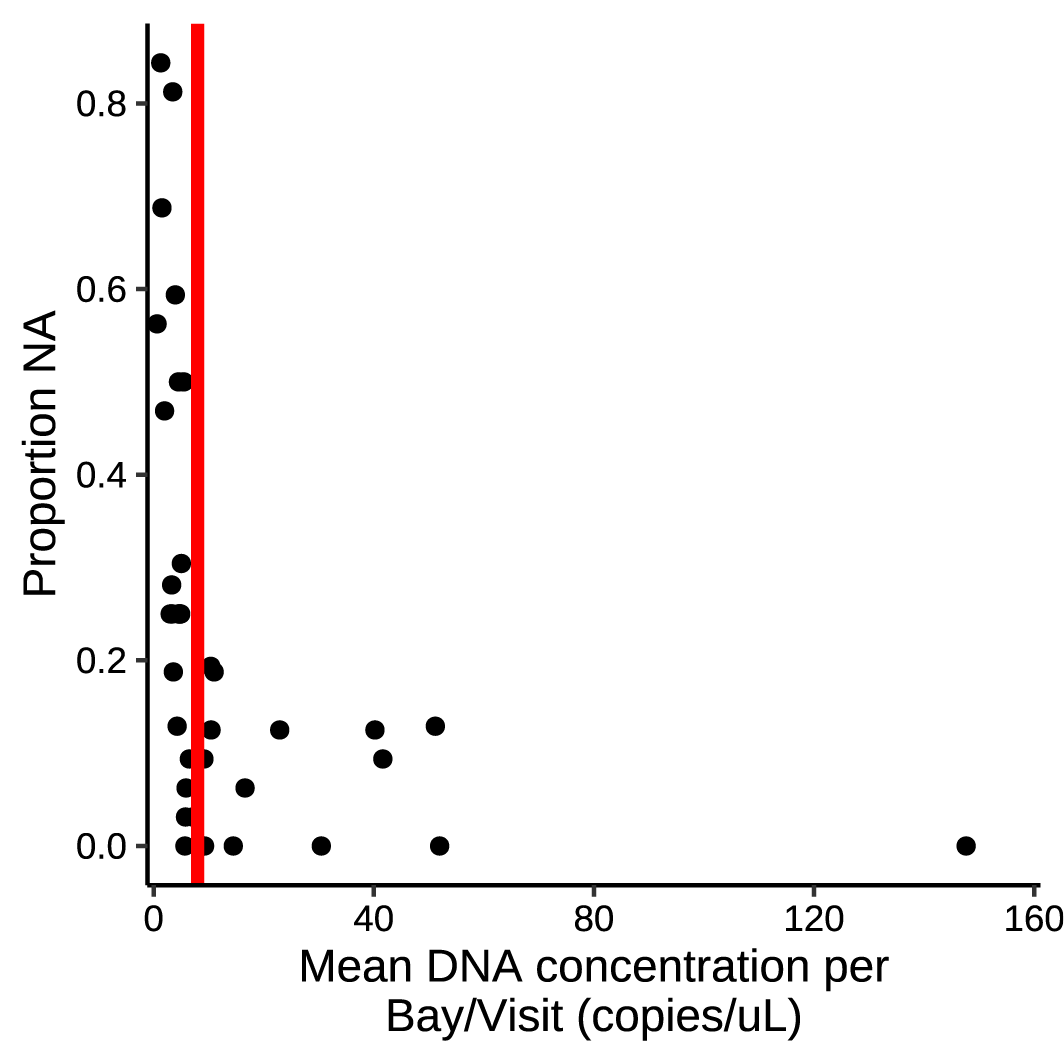

Figure S 4. Proportion of qPCR NA-values as a function of mean eDNA concentration per bay and visit. The vertical red line denotes a subjective threshold concentration for the delineation of true negatives. Non-detects below this line were assigned a values of zero while non-detects above the threshold were left unchanged.

.

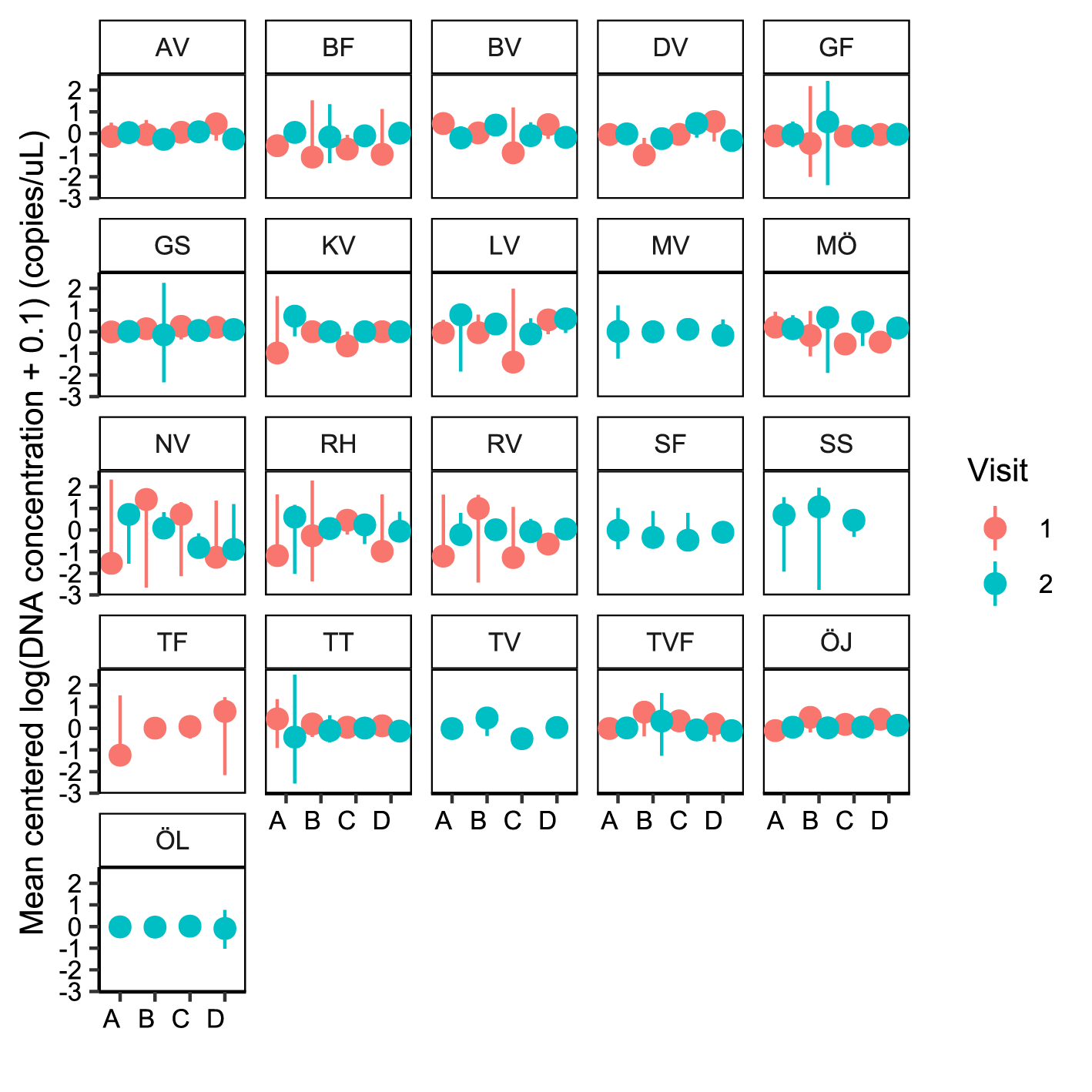

Figure S 5. Transect plot showing mean centered and log transformed eDNA concentrations (copies µL^-1^) per bay, transect (A-D) and visit (1 and 2). Each visit consists of two filter replicates where each filter consists of four technical qPCR replicates. Bay ID: AV = Askviken, BF = Björnöfjärden/Torpe Infjärd, BV = Byviken/Ryssundet, DV = Dalviken, GF = Gisslingöfladen, GS = Granösundet, KV = Kyrkviken/Utö, LV = Lännåkersviken, MV = Myttingeviken, MÖ = Mulö/Lögla, NV = Nynäsviken, RH = Rotholmaviken, RV = Rassa vikar, SF = Söderöfjärden/Sladdarön, SP = Släpan/Ekefjärd, SS = Södersundet, TF = Tofladen/Gropaviken, TT = Tomtviken/Urö, TV = Tranvik/Djuröviken, TVF = Tranviksfjärden, ÖJ = Öjaren/Söderöra/Norröra, ÖL = Östra Lemaren.

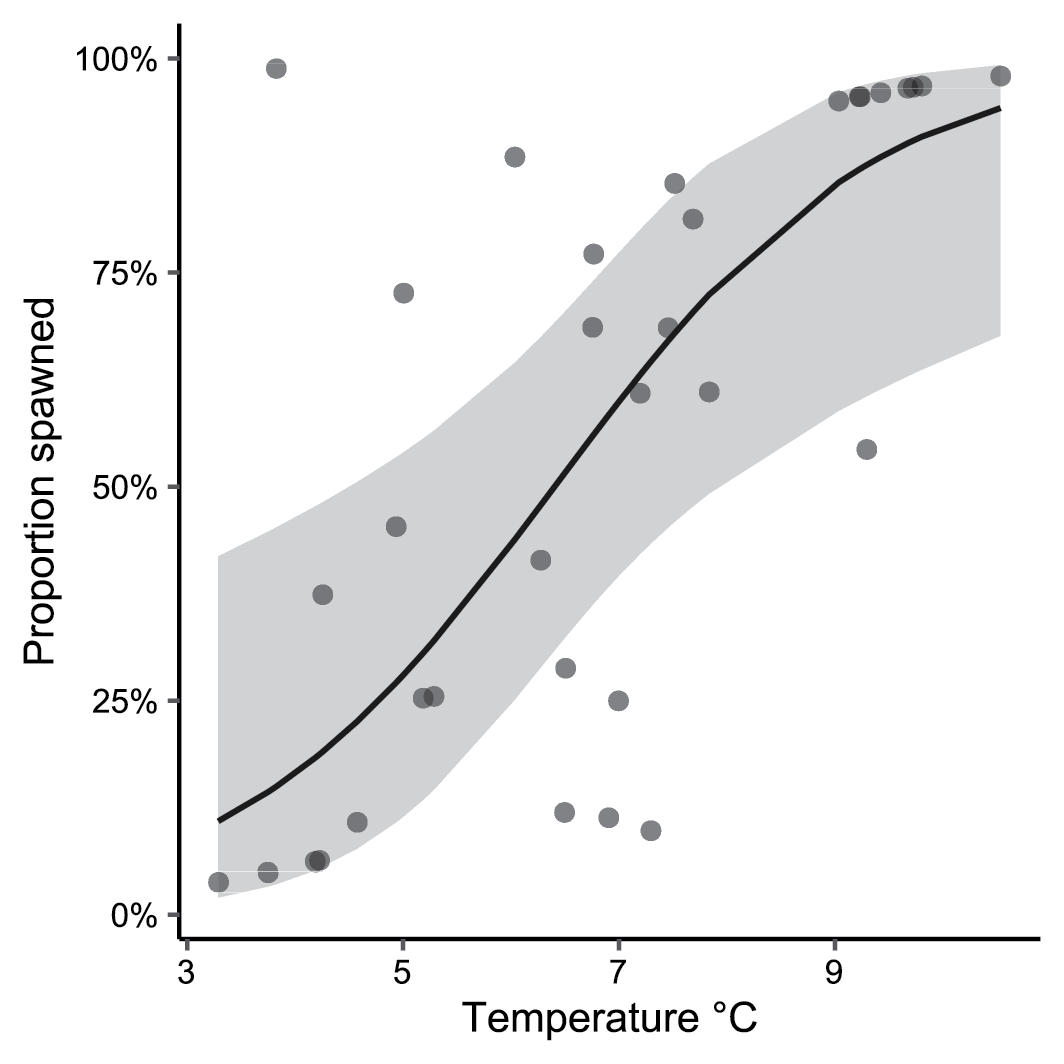

Figure S 6. Proportion of spawned pike per bay and visit as a function of temperature. The relationship was fitted using a GLM with a binomial distribution (estimate 0.67, se=0.24, z=2.76, p=0.0058) and plotted using ggpredict.

**Tables**

Table S 1. Summary statistics (mean ± (SD)) for measured abiotic variables per bay and visit. Bay ID: AV = Askviken, BF = Björnöfjärden/Torpe Infjärd, BV = Byviken/Ryssundet, DV = Dalviken, GF = Gisslingöfladen, GS = Granösundet, KV = Kyrkviken/Utö, LV = Lännåkersviken, MV = Myttingeviken, MÖ = Mulö/Lögla, NV = Nynäsviken, RH = Rotholmaviken, RV = Rassa vikar, SF = Söderöfjärden/Sladdarön, SP = Släpan/Ekefjärd, SS = Södersundet, TF = Tofladen/Gropaviken, TT = Tomtviken/Urö, TV = Tranvik/Djuröviken, TVF = Tranviksfjärden, ÖJ = Öjaren/Söderöra/Norröra, ÖL = Östra Lemaren.

| **Bay ID** | **Visit** | **Depth**  **(m)** | **Salinity**  **(PSU)** | **Oxygen**  **(mg/L)** | **Turbidity**  **(FTU)** | **Chlorophyll A**  **(ppb)** | **Temperature**  **(°C)** |
| --- | --- | --- | --- | --- | --- | --- | --- |
| AV | 1 | 1.5 (0.21) | 6.3 (0.01) | 13.9 (0.09) | 2.9 (0.13) | 1.9 (0.19) | 4.2 (0.13) |
| AV | 2 | 1.4 (0.06) | 6.3 (0.08) | 11.5 (0.26) | 5.0 (1.48) | 2.7 (0.21) | 9.7 (0.51) |
| BF | 1 | 1.0 (0.48) | 5.0 (0.01) | 14.9 (0.20) | 1.1 (0.09) | 4.0 (0.50) | 3.3 (0.19) |
| BF | 2 | 1.5 (0.78) | 5.1 (0.02) | 12.9 (0.14) | 1.2 (0.24) | 2.8 (0.32) | 9.0 (0.51) |
| BV | 1 | 0.8 (0.65) | 5.9 (0.27) | 14.0 (0.31) | 2.1 (0.28) | 2.8 (0.68) | 4.3 (0.47) |
| BV | 2 | 0.4 (0.41) | 6.3 (0.19) | 12.5 (0.44) | 1.6 (0.28) | 1.8 (0.48) | 7.5 (0.92) |
| DV | 1 | 1.8 (0.63) | 5.2 (0.01) | 15.0 (0.19) | 0.9 (0.02) | 1.9 (0.17) | 3.8 (0.23) |
| DV | 2 | 1.6 (0.46) | 5.2 (0.00) | 14.0 (0.18) | 0.6 (0.02) | 1.2 (0.08) | 6.5 (0.39) |
| GF | 1 | 0.7 (0.06) | 5.5 (0.04) | 13.1 (0.40) | 1.5 (1.08) | 4.1 (0.33) | 6.3 (0.35) |
| GF | 2 | 0.3 (0.21) | 5.6 (0.06) | 12.7 (0.61) | 1.7 (0.44) | 2.5 (0.47) | 7.0 (0.86) |
| GS | 1 | 0.5 (0.09) | 5.9 (0.01) | 13.8 (0.44) | 1.2 (0.43) | 1.0 (0.04) | 7.3 (0.75) |
| GS | 2 | 0.5 (0.40) | 5.8 (0.01) | 12.7 (0.15) | 1.7 (1.35) | 0.9 (0.03) | 7.7 (0.43) |
| KV | 1 | 1.0 (0.58) | 6.7 (0.01) | 15.1 (0.24) | 0.8 (0.08) | 1.0 (0.21) | 3.9 (0.14) |
| KV | 2 | 1.2 (0.48) | 6.6 (0.03) | 14.0 (0.45) | 0.9 (0.02) | 0.8 (0.07) | 6.0 (0.25) |
| LV | 1 | 1.2 (0.44) | 6.1 (0.06) | 13.8 (0.15) | 6.0 (5.51) | 2.8 (0.31) | 4.9 (0.10) |
| LV | 2 | 0.6 (0.29) | 6.4 (0.01) | 11.6 (0.08) | 4.4 (2.02) | 2.1 (0.09) | 9.2 (0.07) |
| MV | 2 | 1.7 (0.15) | 2.6 (0.31) | 15.6 (0.42) | 1.5 (0.19) | 5.9 (1.26) | 8.6 (1.29) |
| MÖ | 1 | 1.4 (0.41) | 5.4 (0.00) | 13.7 (0.11) | 0.7 (0.15) | 1.2 (0.04) | 5.2 (0.12) |
| MÖ | 2 | 1.2 (0.32) | 5.5 (0.00) | 11.3 (0.50) | 3.8 (1.90) | 1.6 (0.20) | 9.8 (0.13) |
| NV | 1 | 1.5 (0.36) | 5.7 (0.07) | 15.1 (0.21) | 3.4 (0.24) | 3.5 (0.57) | 3.7 (0.04) |
| NV | 2 | 1.5 (0.16) | 5.8 (0.09) | 13.6 (0.08) | 2.1 (0.29) | 2.3 (0.17) | 6.8 (0.06) |
| RH | 1 | 1.0 (0.19) | 5.1 (0.00) | 14.3 (0.14) | 1.5 (0.46) | 1.8 (0.07) | 5.0 (0.37) |
| RH | 2 | 1.1 (0.23) | 5.1 (0.00) | 12.7 (0.14) | 2.1 (0.46) | 1.4 (0.06) | 9.2 (0.06) |
| RV | 1 | 1.4 (0.04) | 6.3 (0.02) | 13.5 (0.11) | 1.1 (0.10) | 1.7 (0.25) | 4.2 (0.04) |
| RV | 2 | 1.7 (0.09) | 6.5 (0.01) | 12.8 (0.15) | 0.8 (0.05) | 1.2 (0.21) | 7.2 (0.07) |
| SF | 2 | 1.0 (0.21) | 5.4 (0.03) | 13.1 (0.29) | 1.3 (0.19) | 1.2 (0.11) | 7.8 (0.60) |
| SP | 2 | 0.9 (0.23) | 2.4 (0.17) | 14.8 (0.76) | 5.2 (0.87) | 13.9 (3.43) | 10.4 (0.55) |
| SS | 2 | 0.9 (0.13) | 5.8 (0.00) | 13.7 (0.15) | 0.7 (0.04) | 1.1 (0.03) | 7.5 (0.59) |
| TF | 1 | 1.0 (0.36) | 5.7 (0.02) | 14.1 (0.52) | 0.9 (0.32) | 1.8 (0.24) | 5.3 (0.66) |
| TT | 1 | 0.9 (0.17) | 4.6 (0.08) | 13.9 (0.32) | 2.8 (0.39) | 2.9 (0.13) | 6.5 (0.30) |

*Table S 1 continued.*

| **Bay ID** | **Visit** | **Depth**  **(m)** | **Salinity**  **(PSU)** | **Oxygen**  **(mg/L)** | **Turbidity**  **(FTU)** | **Chlorophyll A**  **(ppb)** | **Temperature**  **(°C)** |
| --- | --- | --- | --- | --- | --- | --- | --- |
| TT | 2 | 1.3 (0.26) | 5.0 (0.03) | 12.6 (0.14) | 2.1 (0.11) | 2.6 (0.23) | 9.7 (0.21) |
| TV | 2 | 1.0 (0.34) | 4.3 (0.01) | 12.6 (0.28) | 2.0 (0.84) | 2.8 (0.26) | 10.5 (0.33) |
| TVF | 1 | 0.4 (0.08) | 5.2 (0.04) | 14.3 (0.09) | 1.1 (0.33) | 1.9 (0.08) | 4.6 (0.09) |
| TVF | 2 | 0.6 (0.19) | 5.4 (0.00) | 13.2 (0.21) | 3.7 (0.82) | 2.3 (0.31) | 9.3 (0.11) |
| ÖJ | 1 | 0.6 (0.12) | 5.9 (0.01) | 13.6 (0.27) | 0.6 (0.09) | 0.9 (0.02) | 6.9 (0.05) |
| ÖJ | 2 | 0.5 (0.06) | 5.8 (0.01) | 13.0 (0.13) | 0.7 (0.09) | 0.9 (0.05) | 6.8 (0.08) |
| ÖL | 2 | 0.7 (0.10) | 5.8 (0.02) | 13.3 (0.56) | 1.5 (0.51) | 1.3 (0.17) | 9.4 (0.15) |
