## Supplementary maps for "Temperature moderates eDNA-biomass relationships in northern pike"

### Rassa Vikar (RV) 45 ha

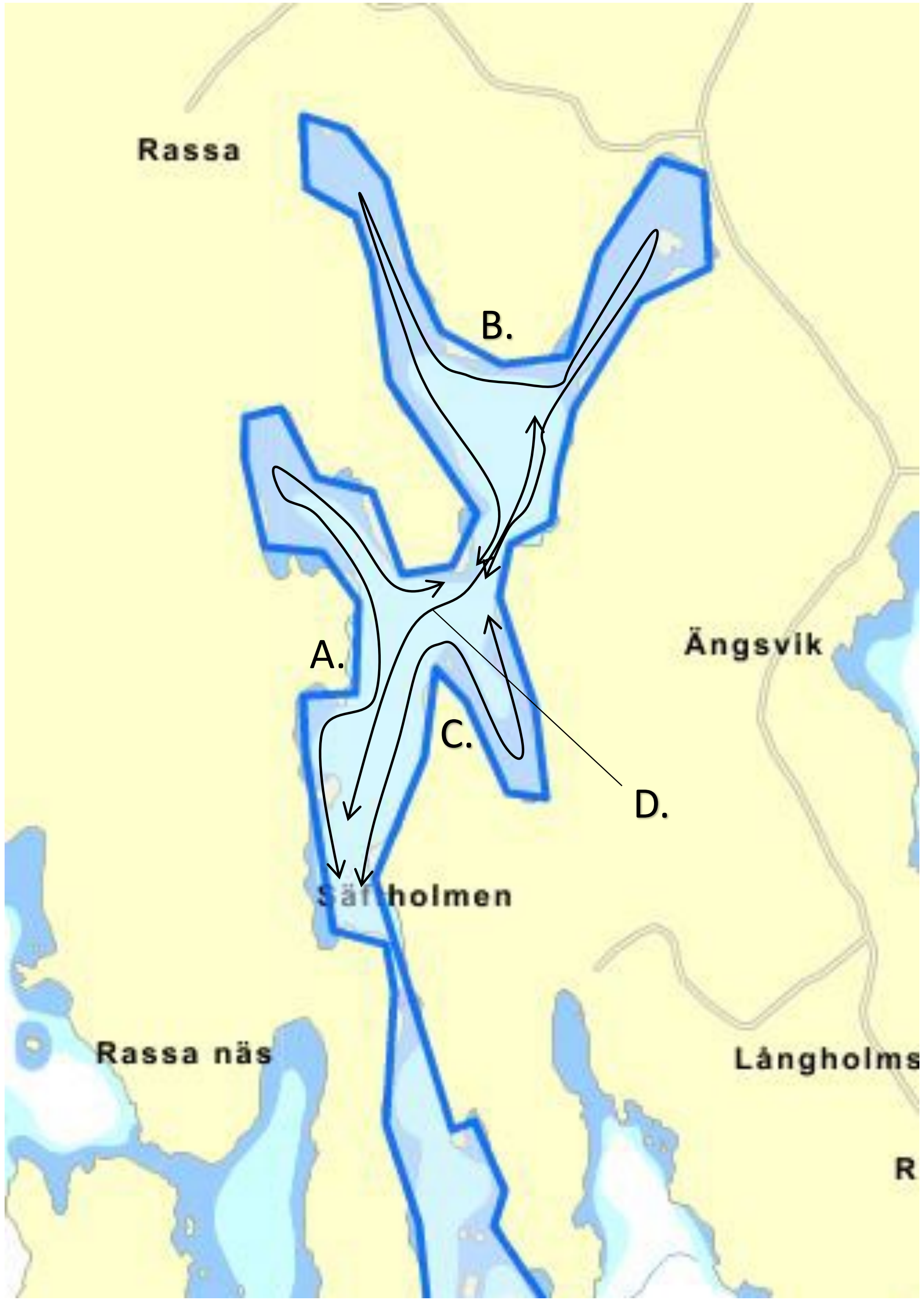

### Nynäsviken (NV) 45 ha

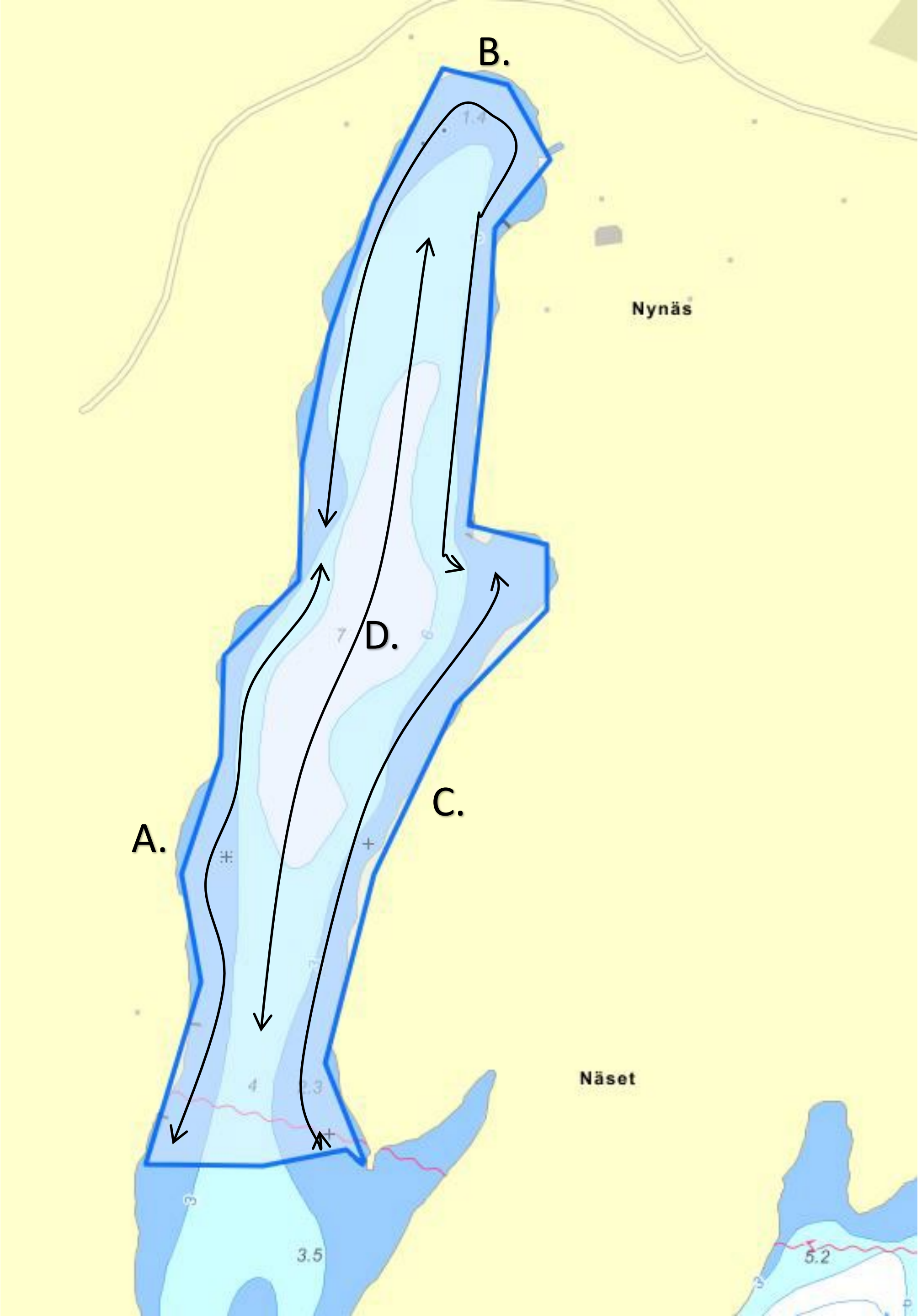

### Byviken/Ryssundet (BV)
# 33 ha

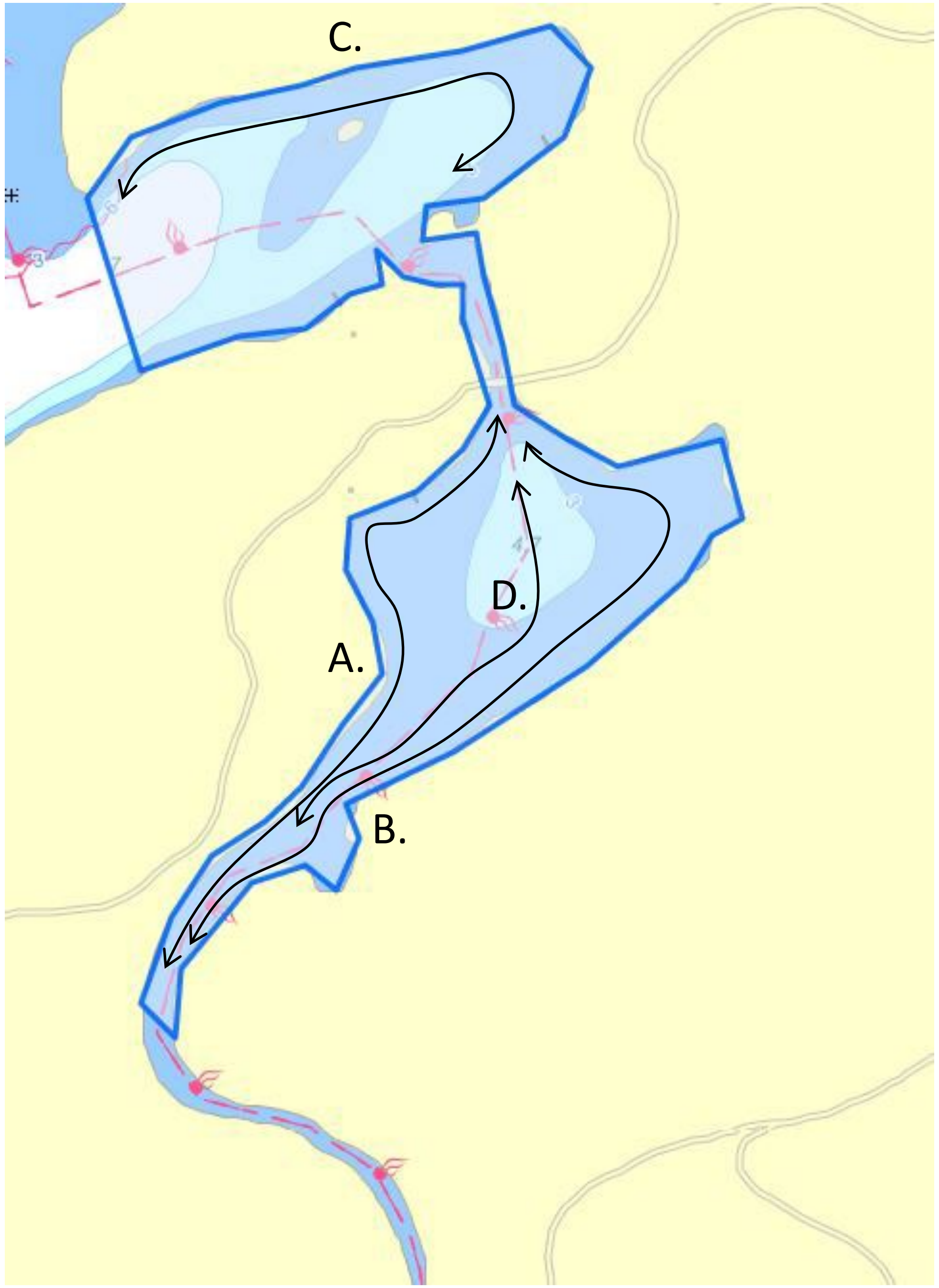

### Kyrkviken/Utö (KV)
# 35 ha

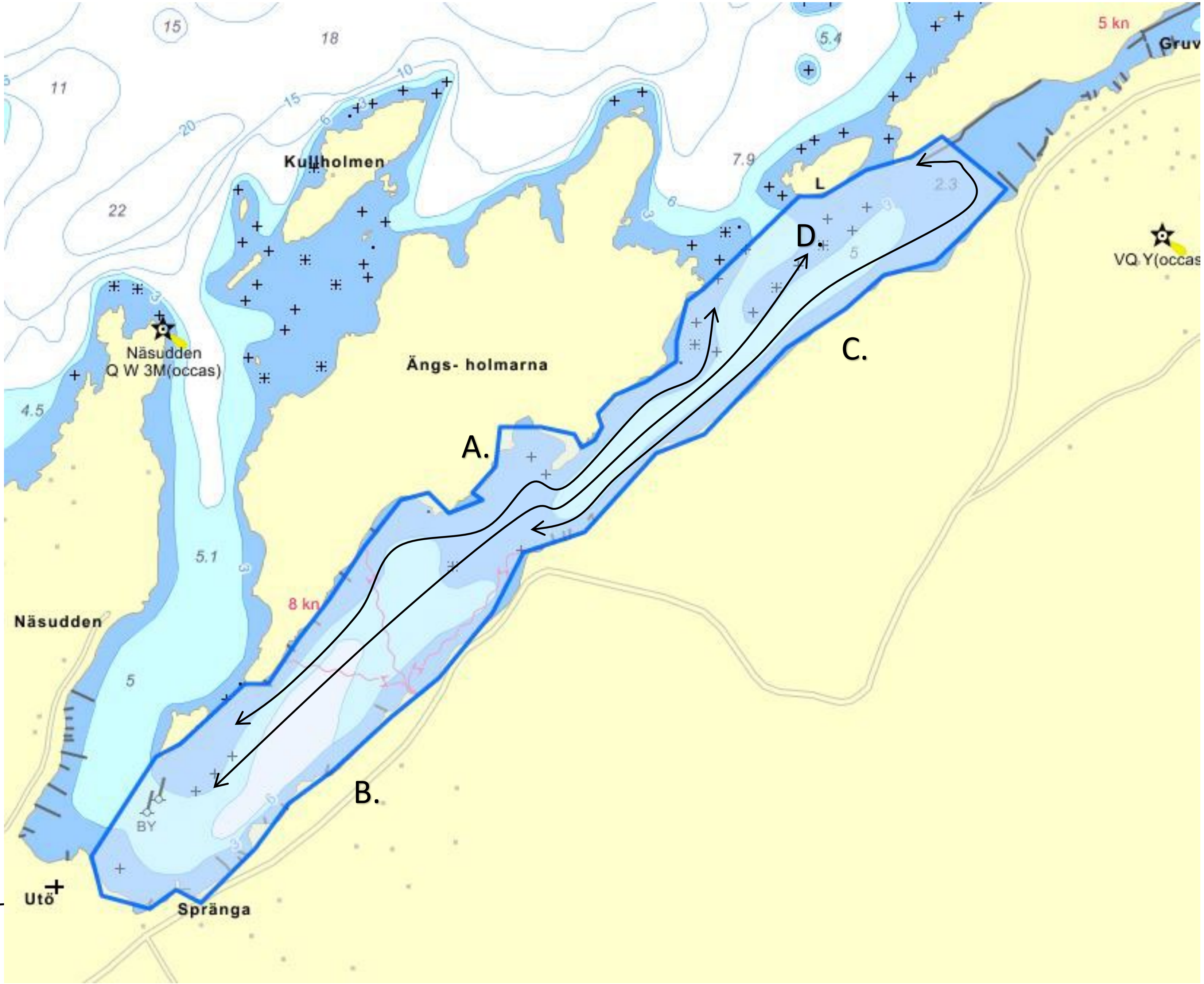

### Björnöfjärden/ Torpe infjärd (BF) 26 ha

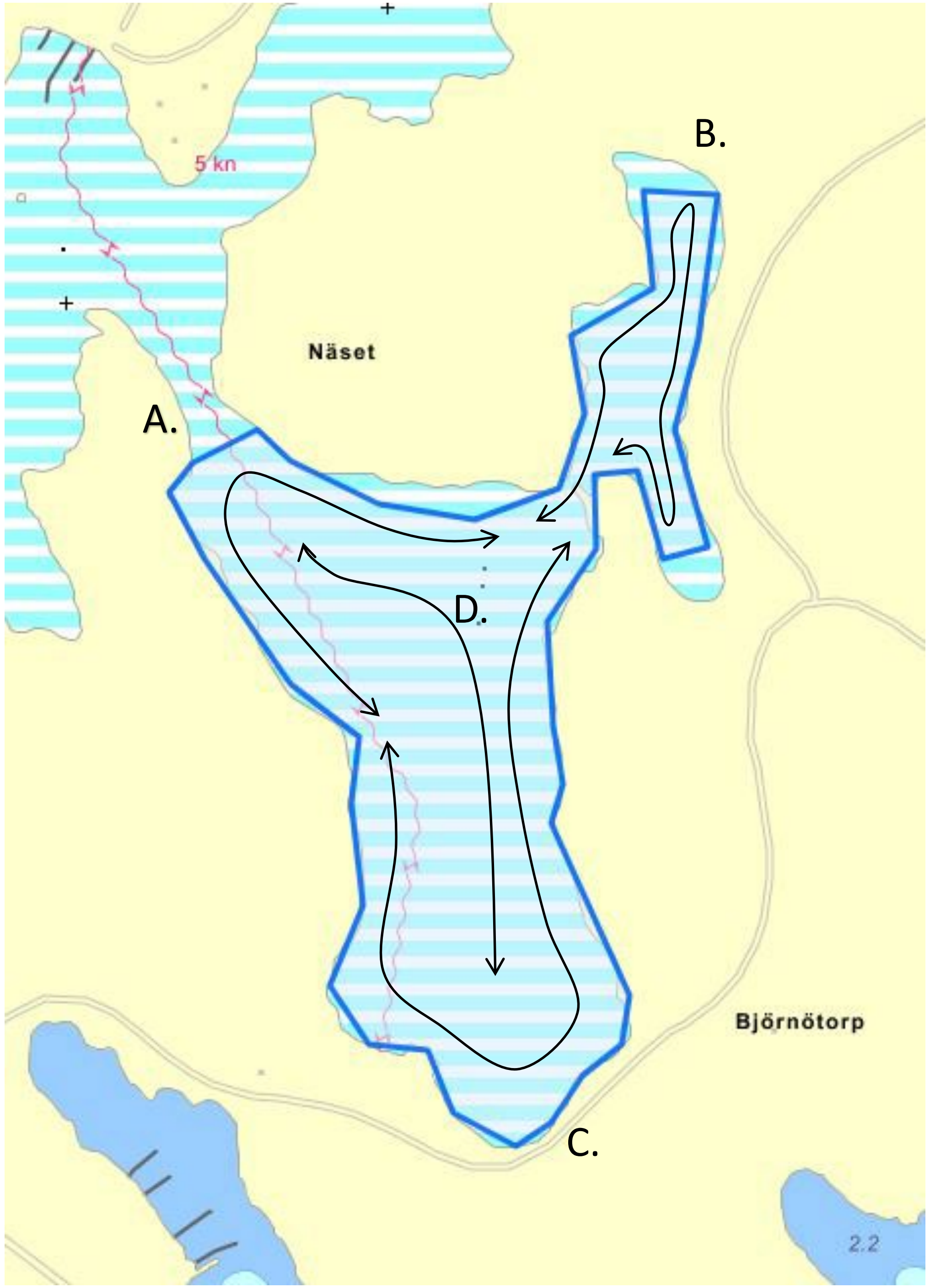

### Tranvik/Djuröviken (TV)
21 ha

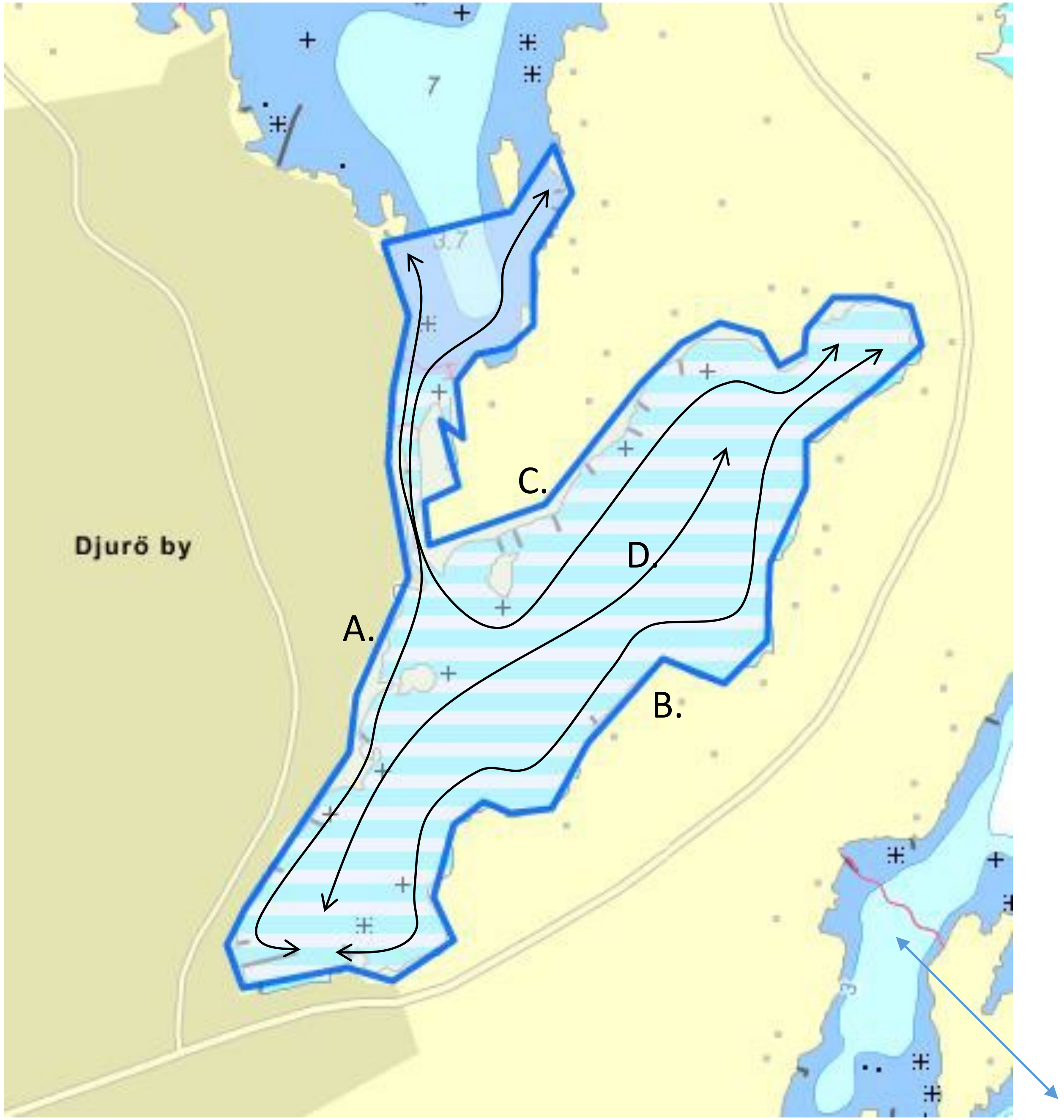

### Tranviksfjärden (TVF)
# 16 ha

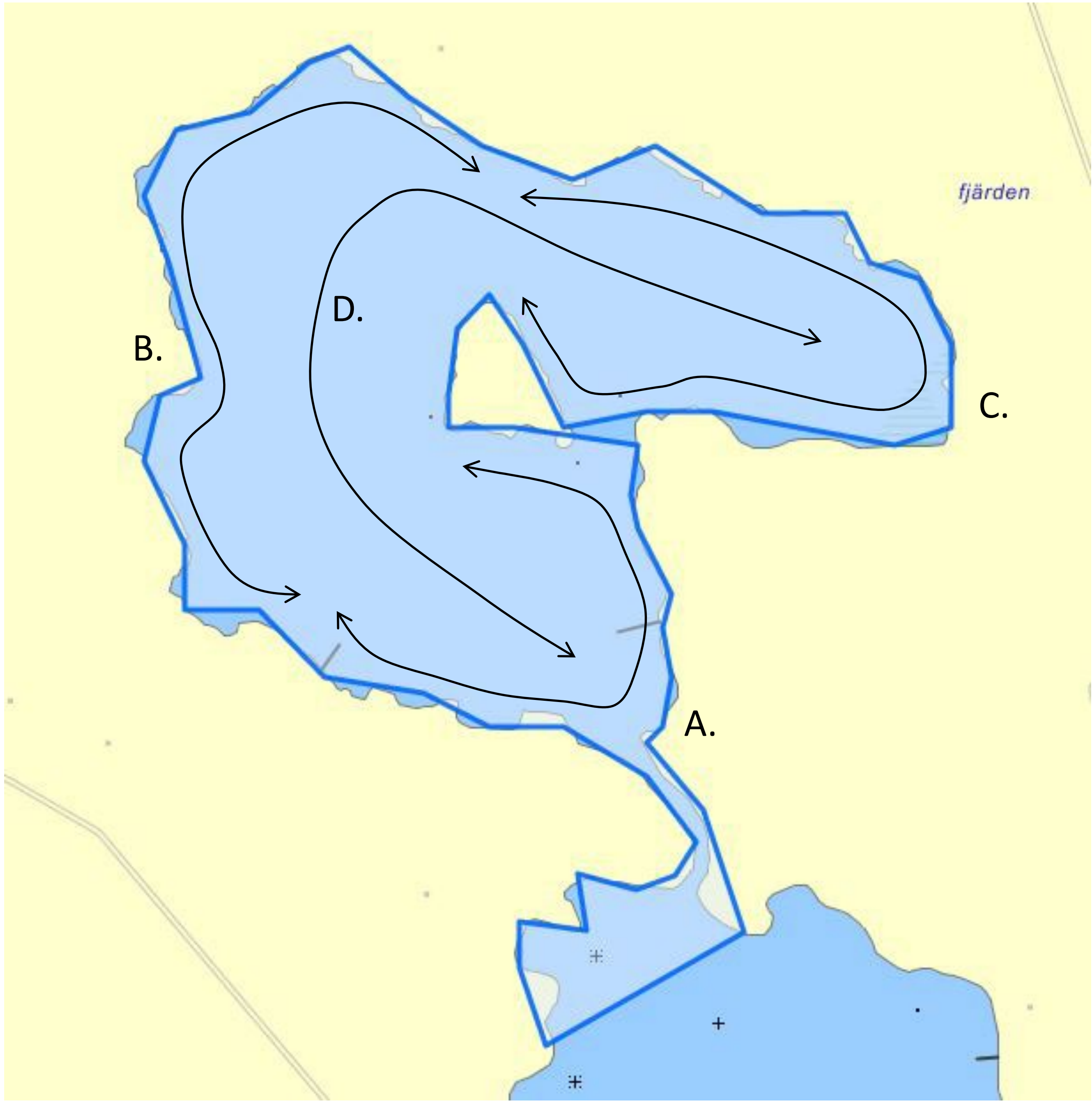

### Söderöfjärden/Sladdarön
# (SF)
# 17 ha

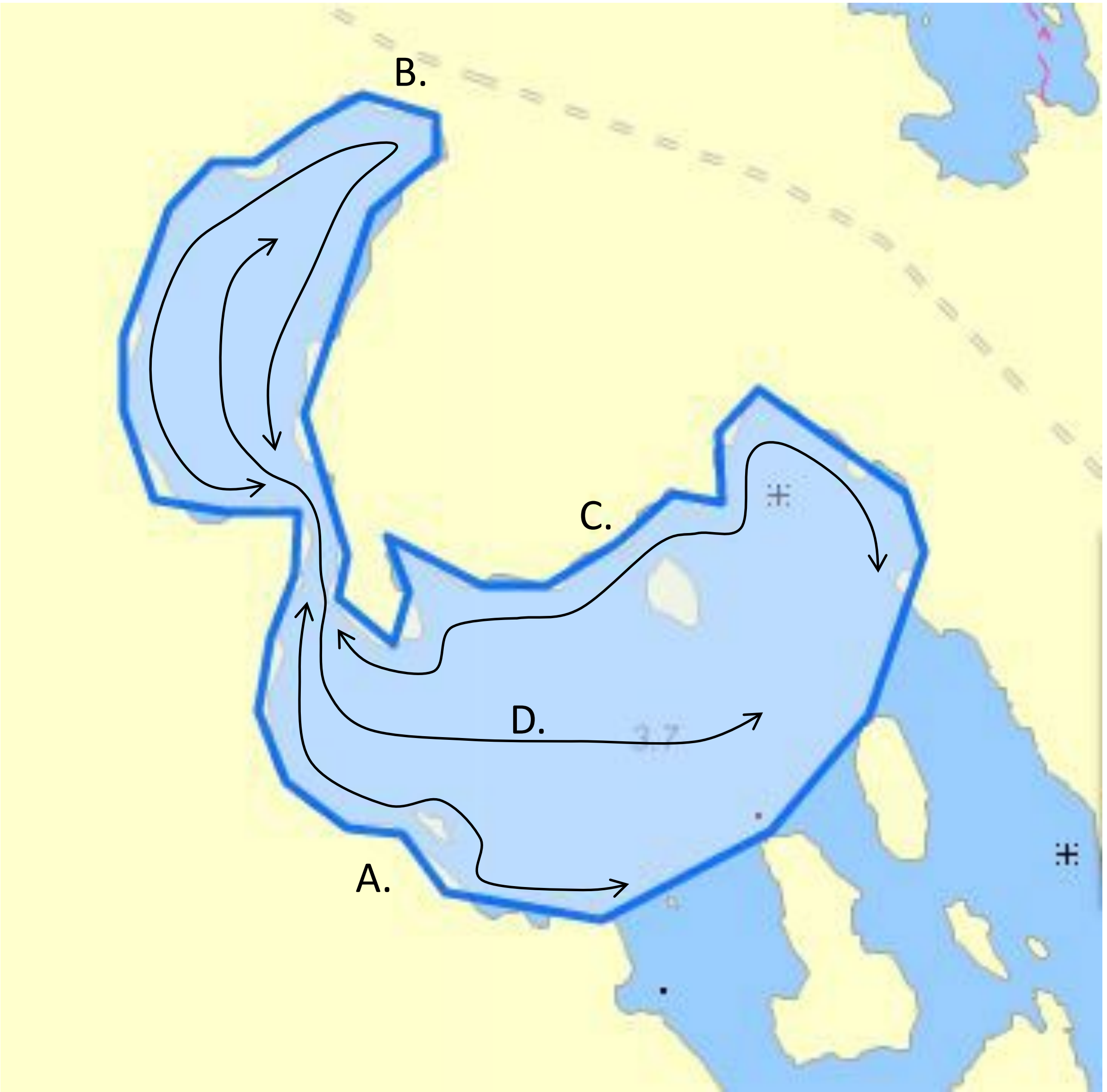

### Släpan/Ekefjärd (SP)
# 43 ha

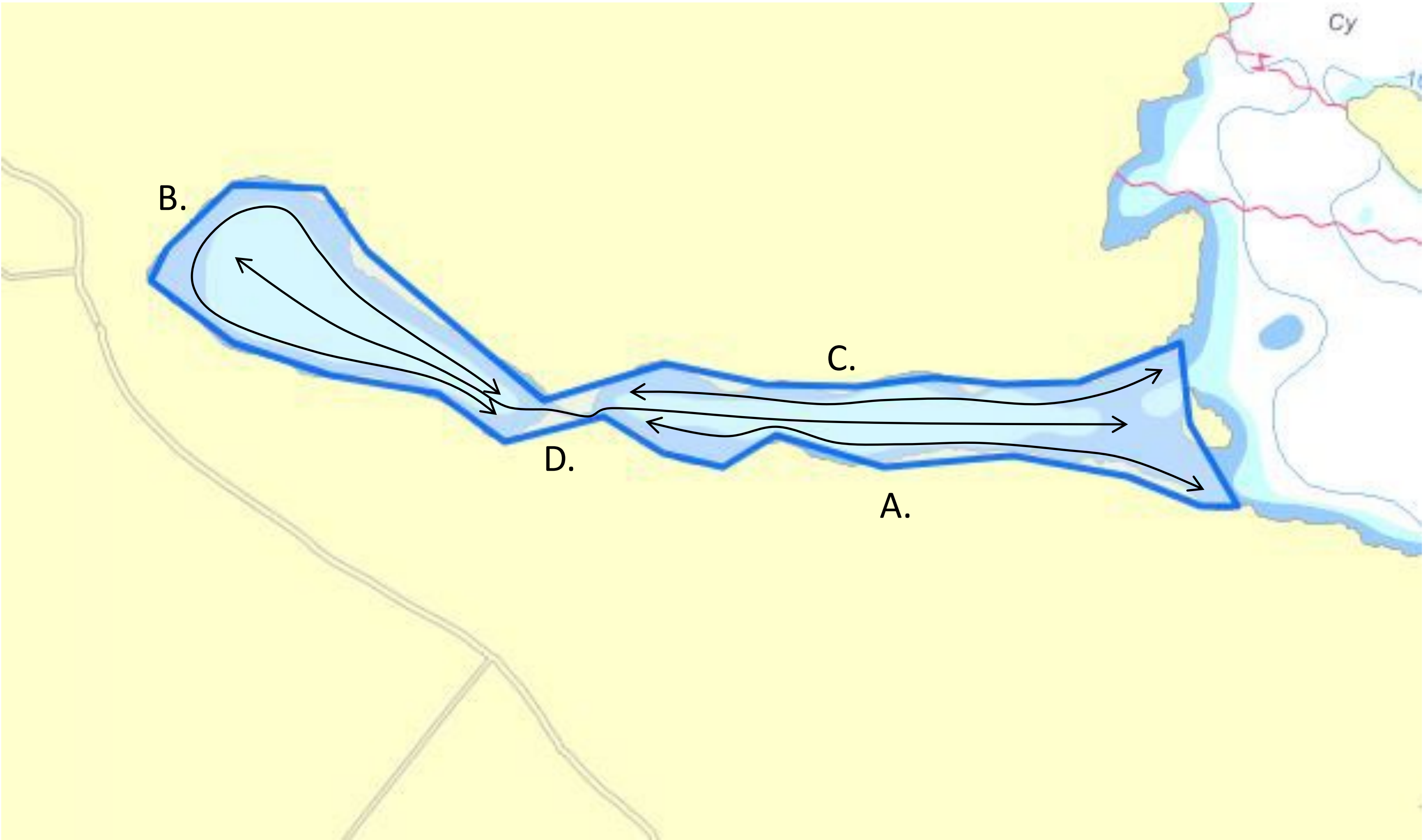

### Myttingeviken (MV)
# 35 ha

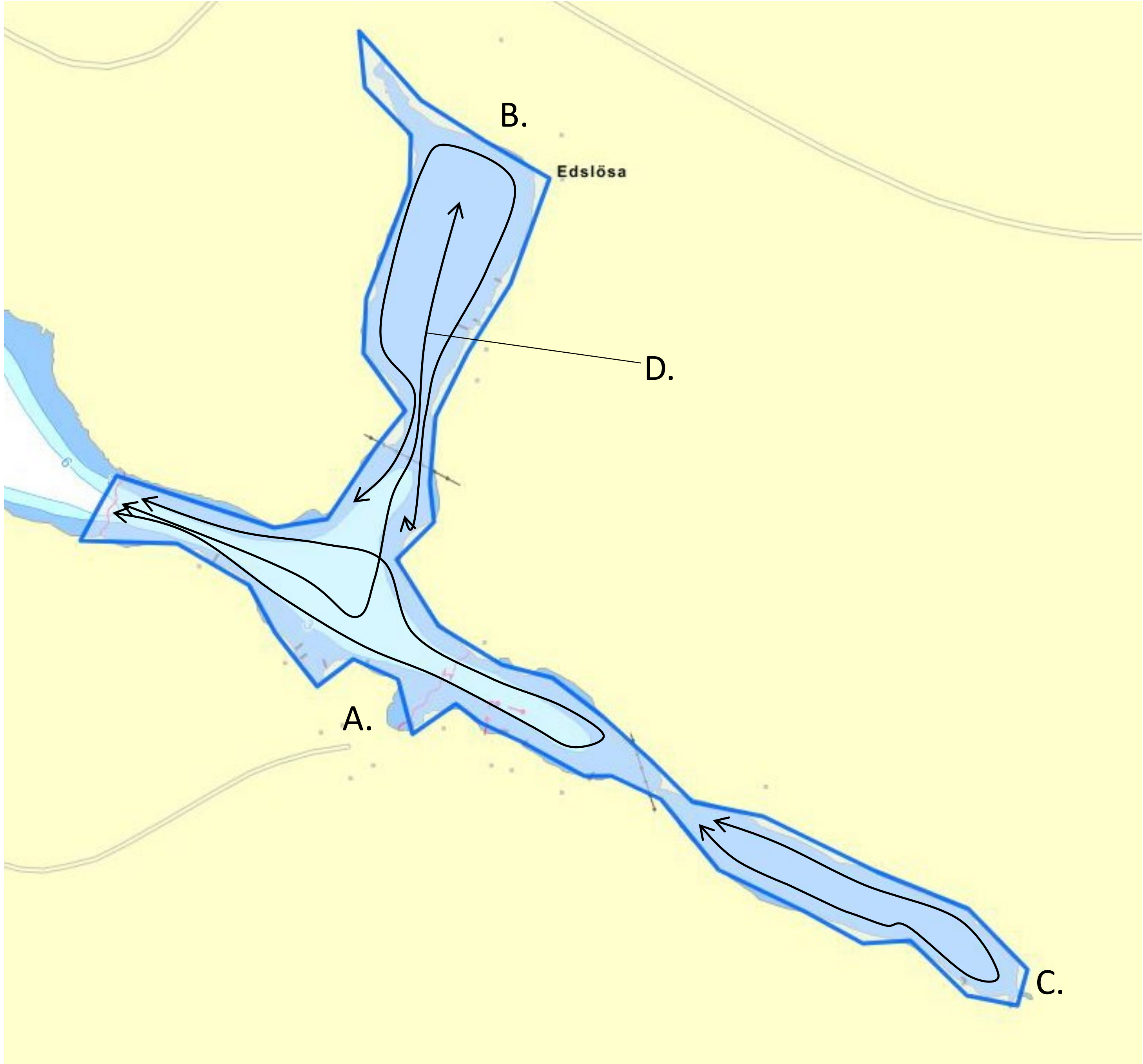

### Lännåkersviken (LV) 45 ha

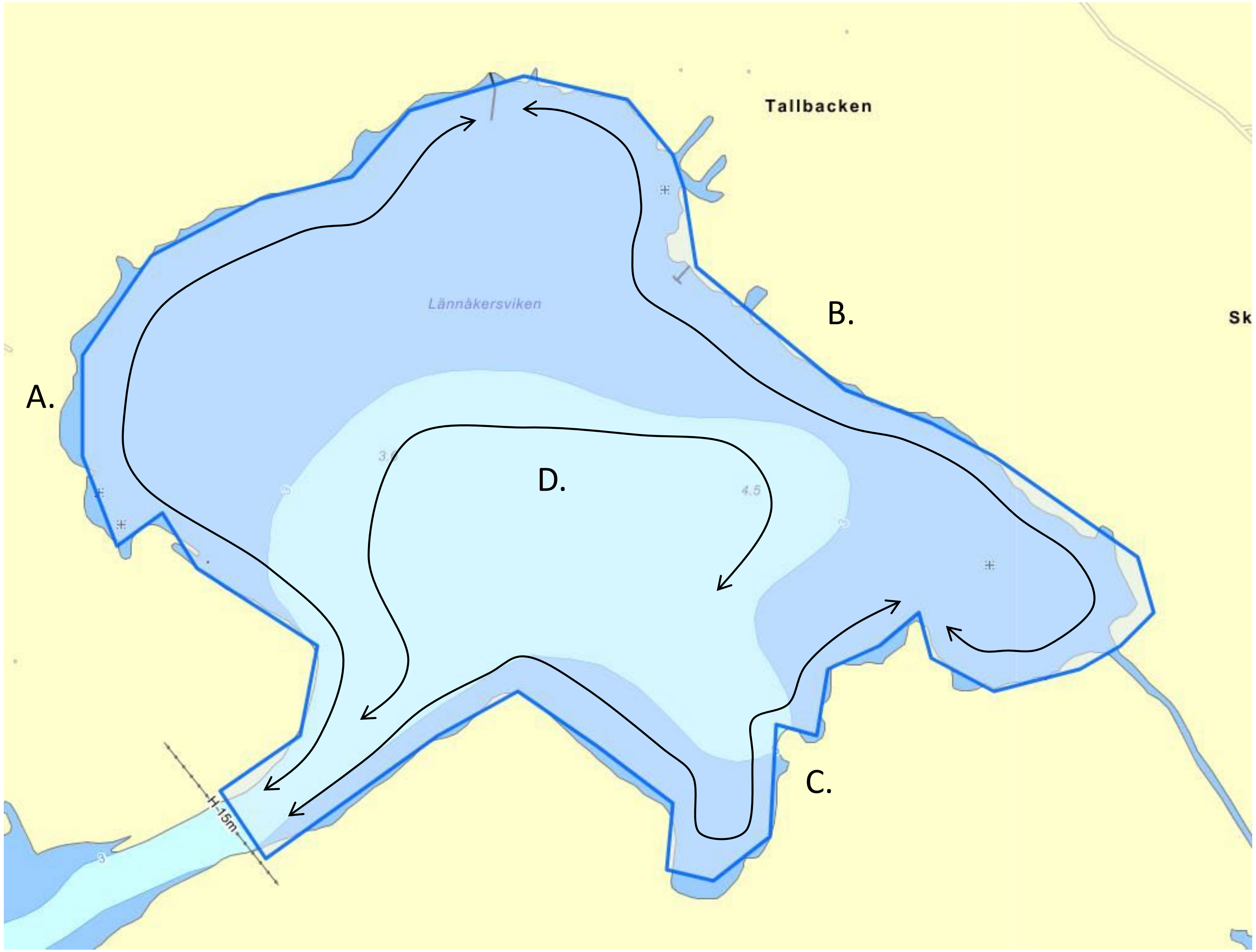

### Askviken (AV) 40 ha

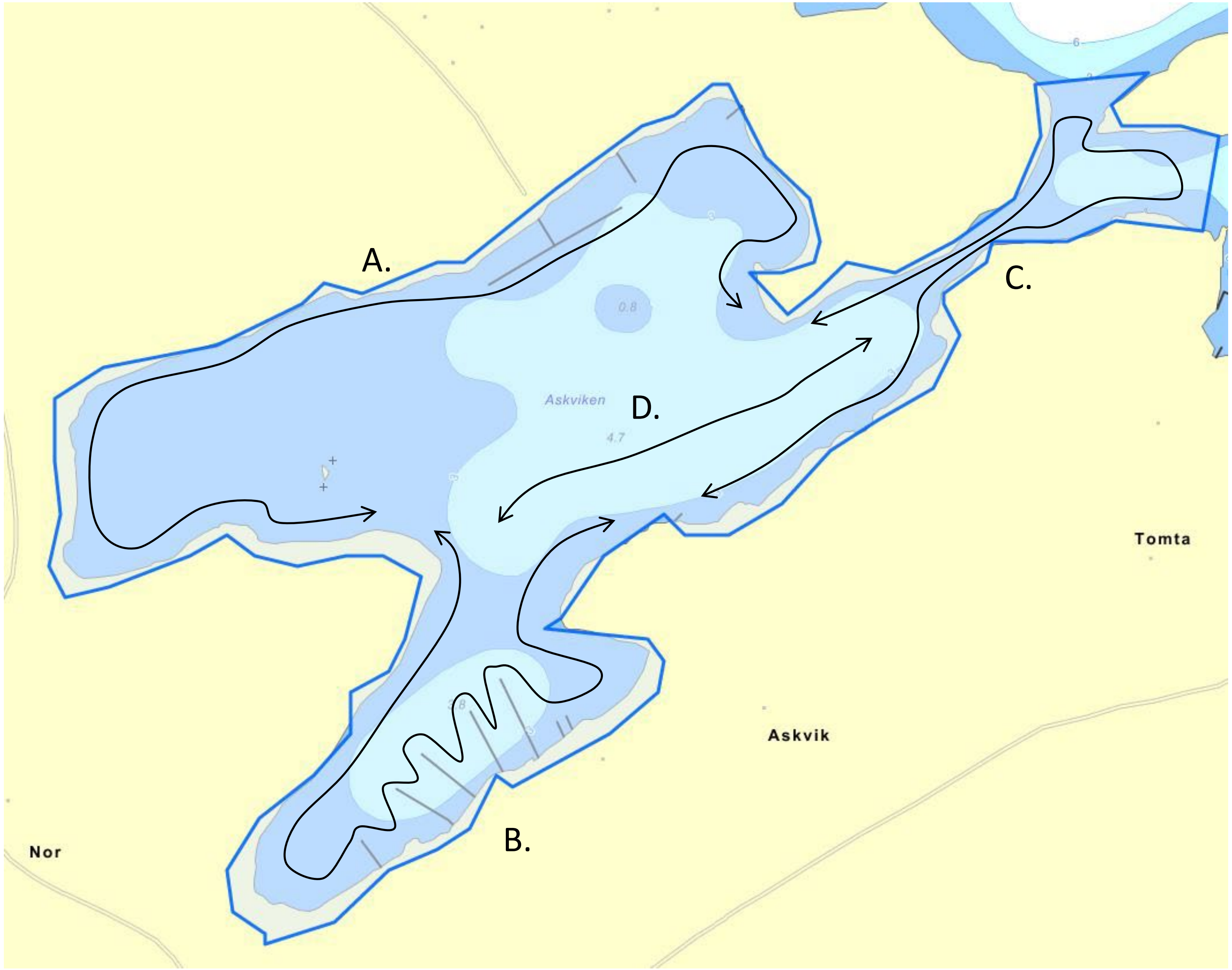

### Dalviken (DV) 36 ha

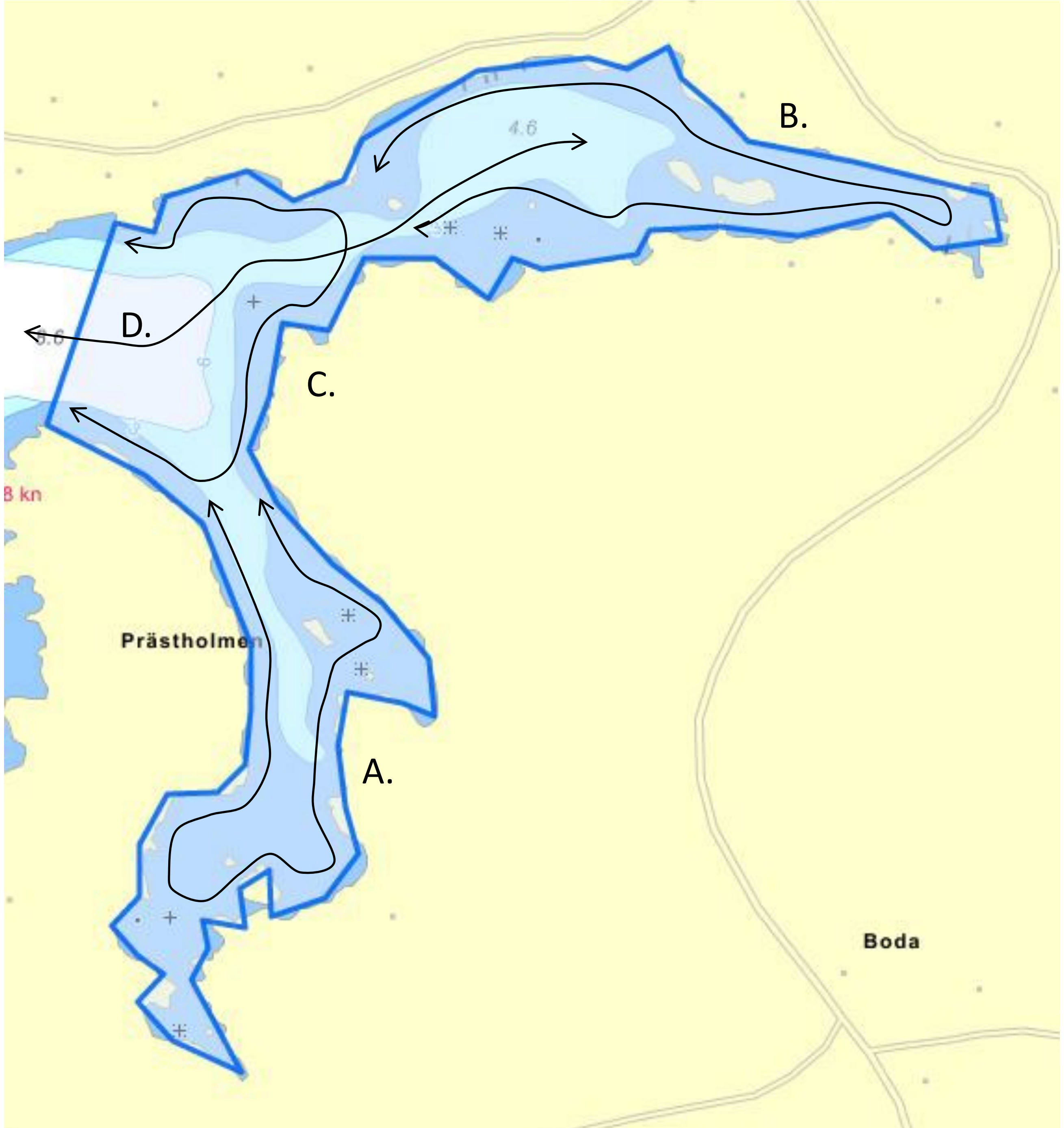

### Rotholmaviken (RH)
# 40 ha

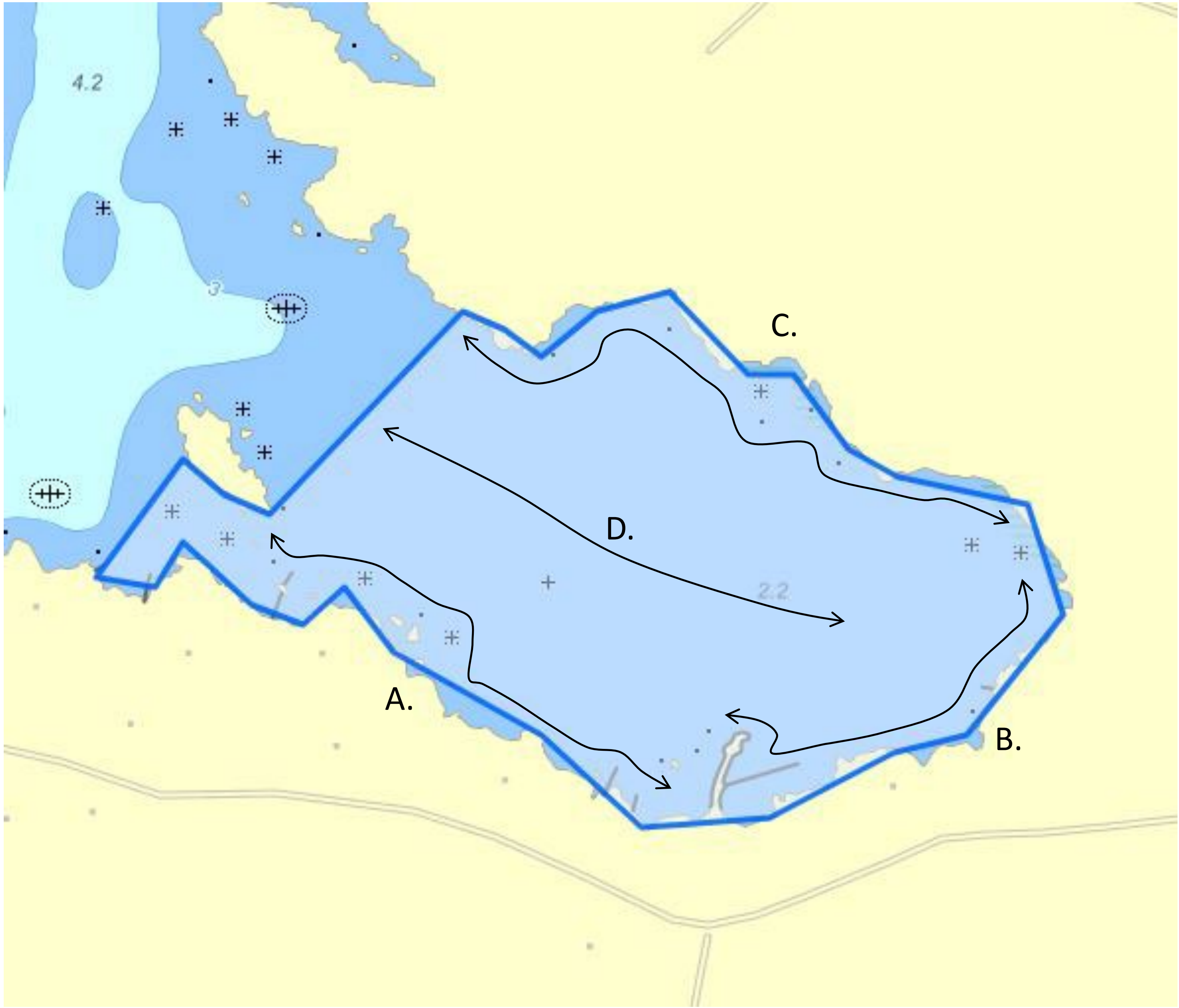

### Gisslingöfladen (GF)
# 13 ha

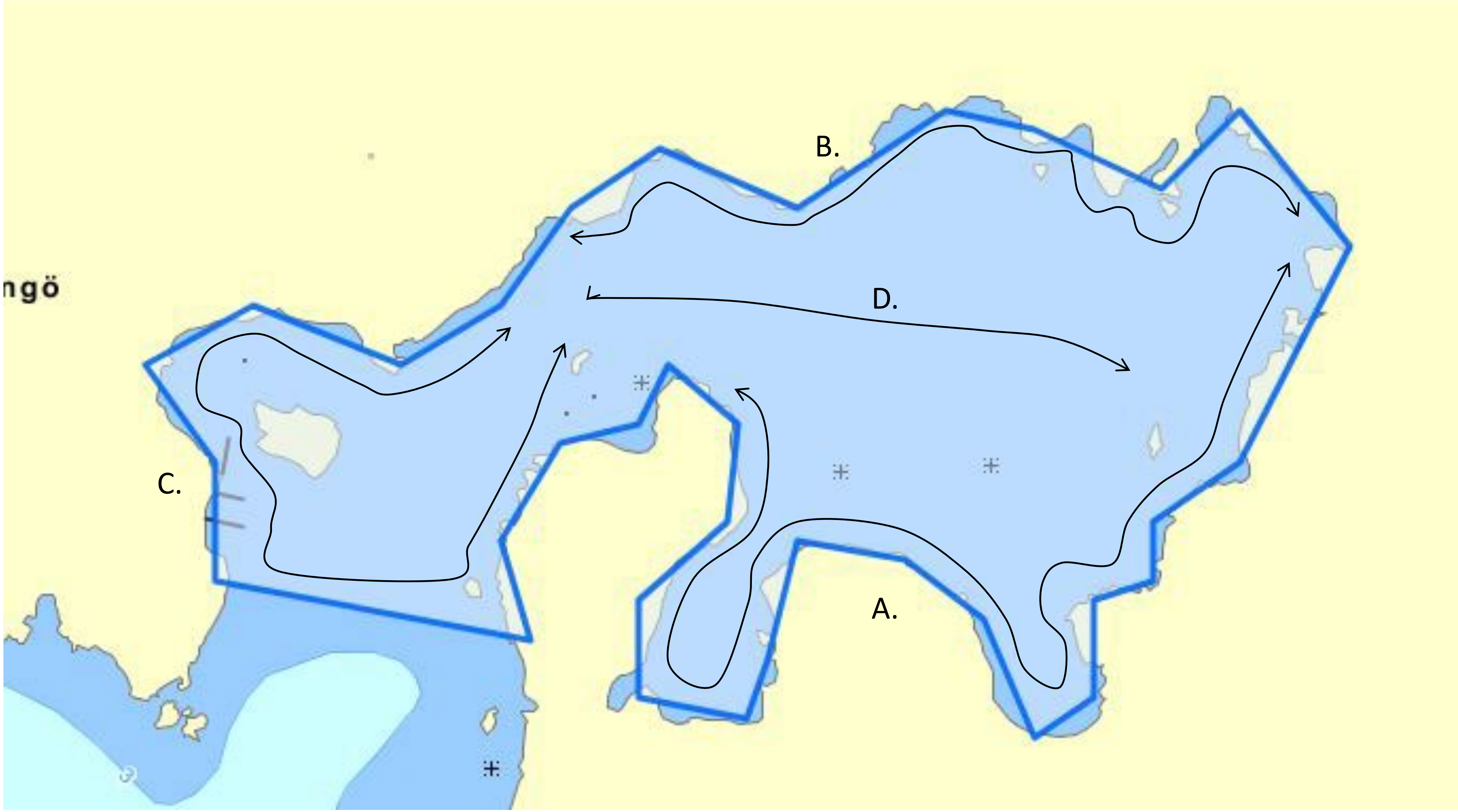

### Tofladen/ Gropaviken (TF) 13 ha

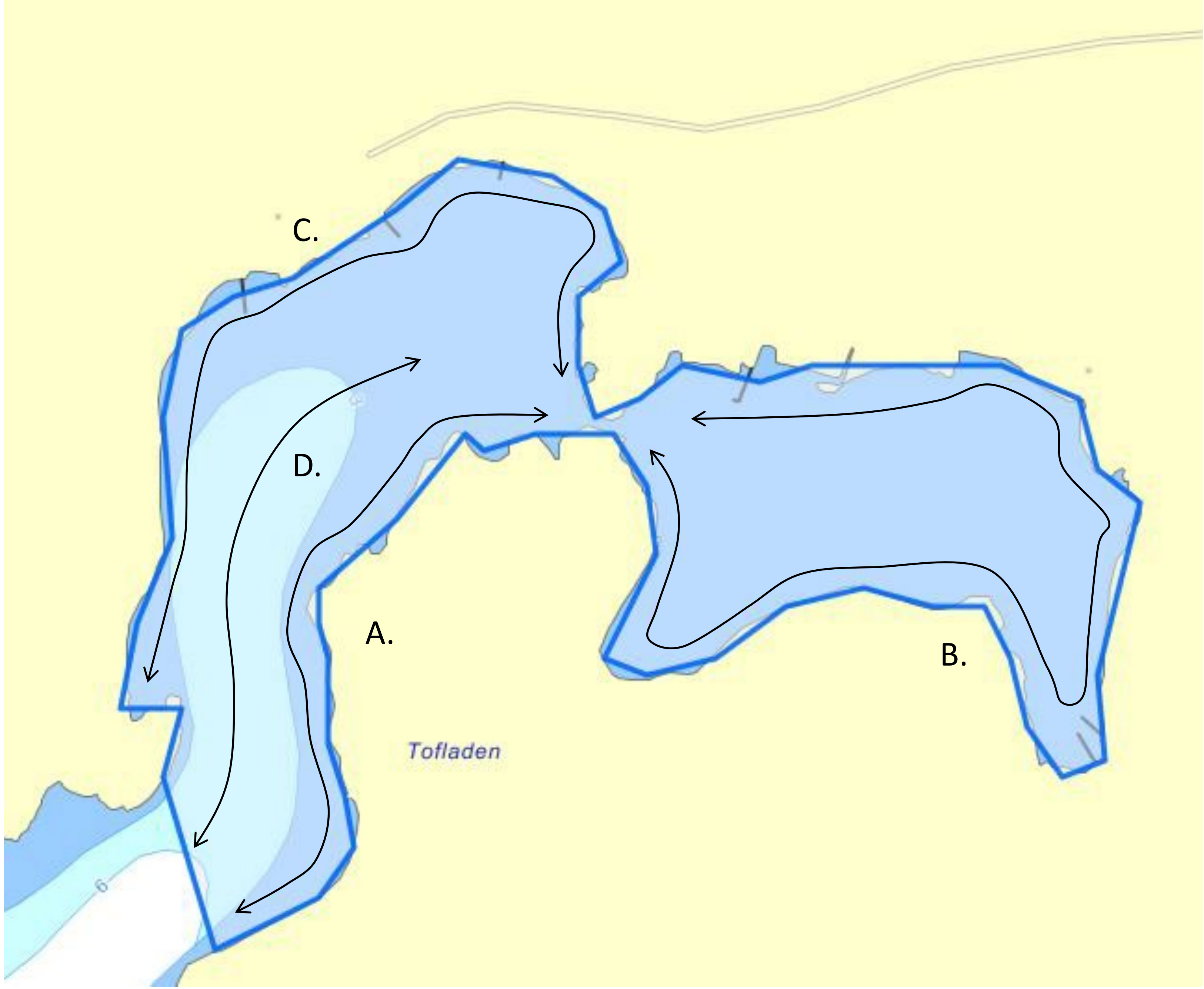

### Södersundet
# (SS)
# 12 ha

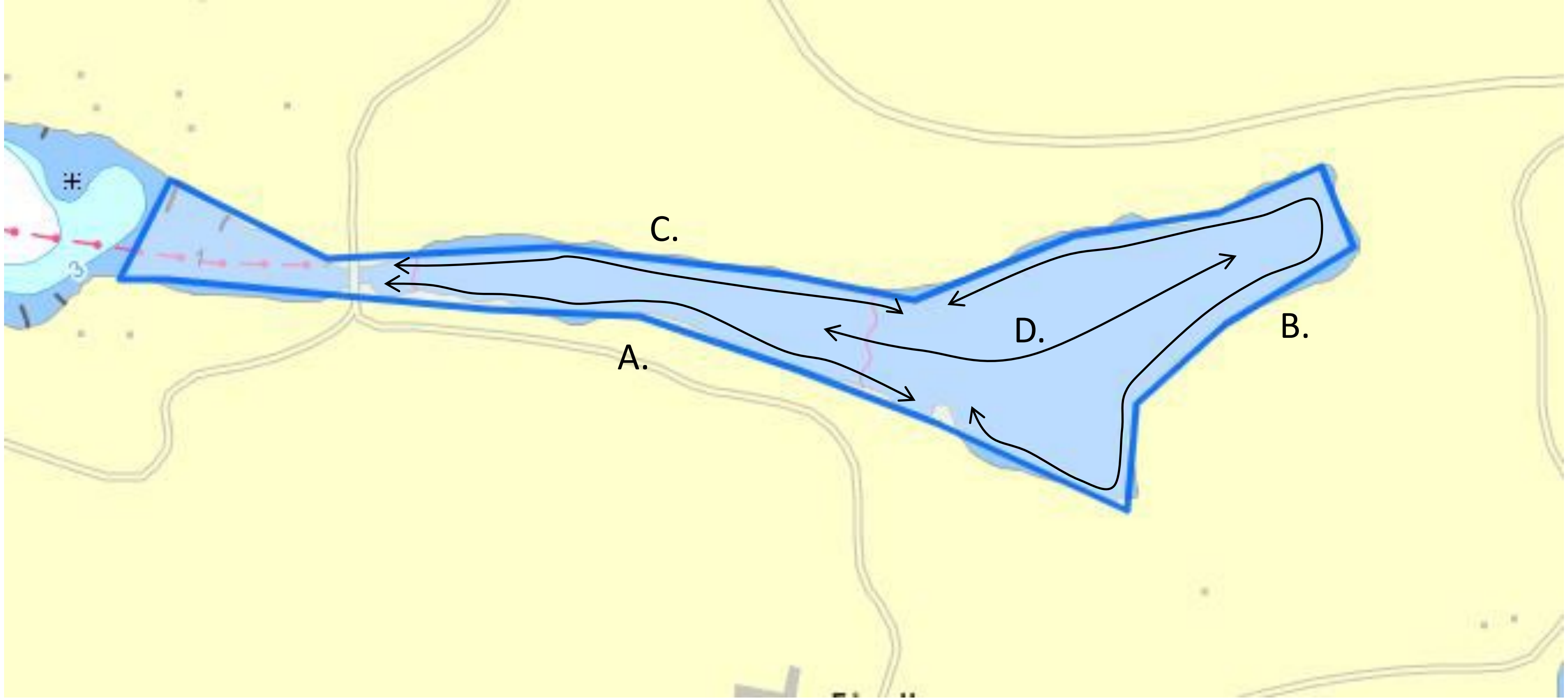

### Östra Lemaren (ÖL) 12 ha

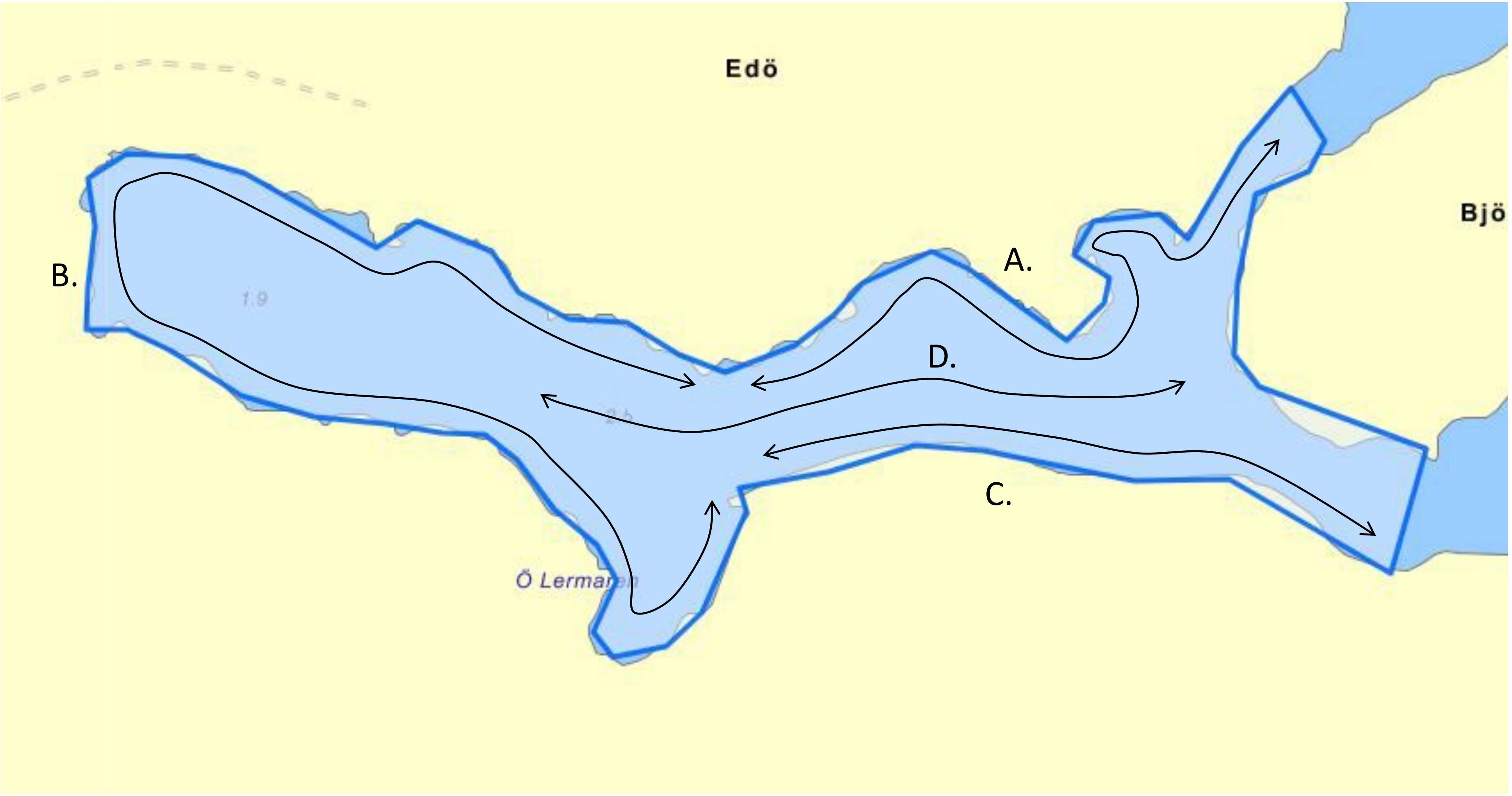

### Öjaren/ Söderöra/ Norröra (ÖJ) 25 ha

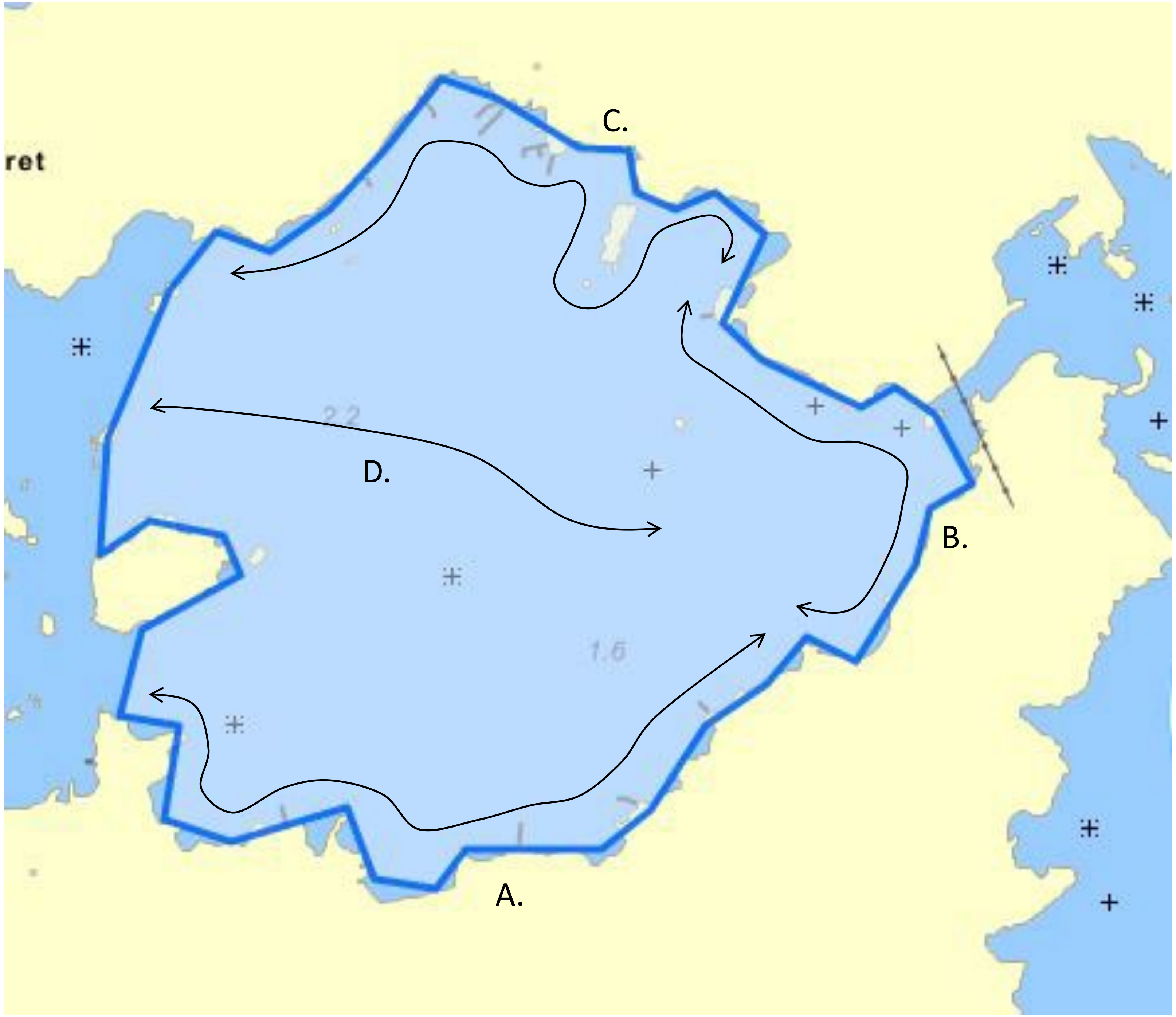

### Granösundet (GS) 40 ha

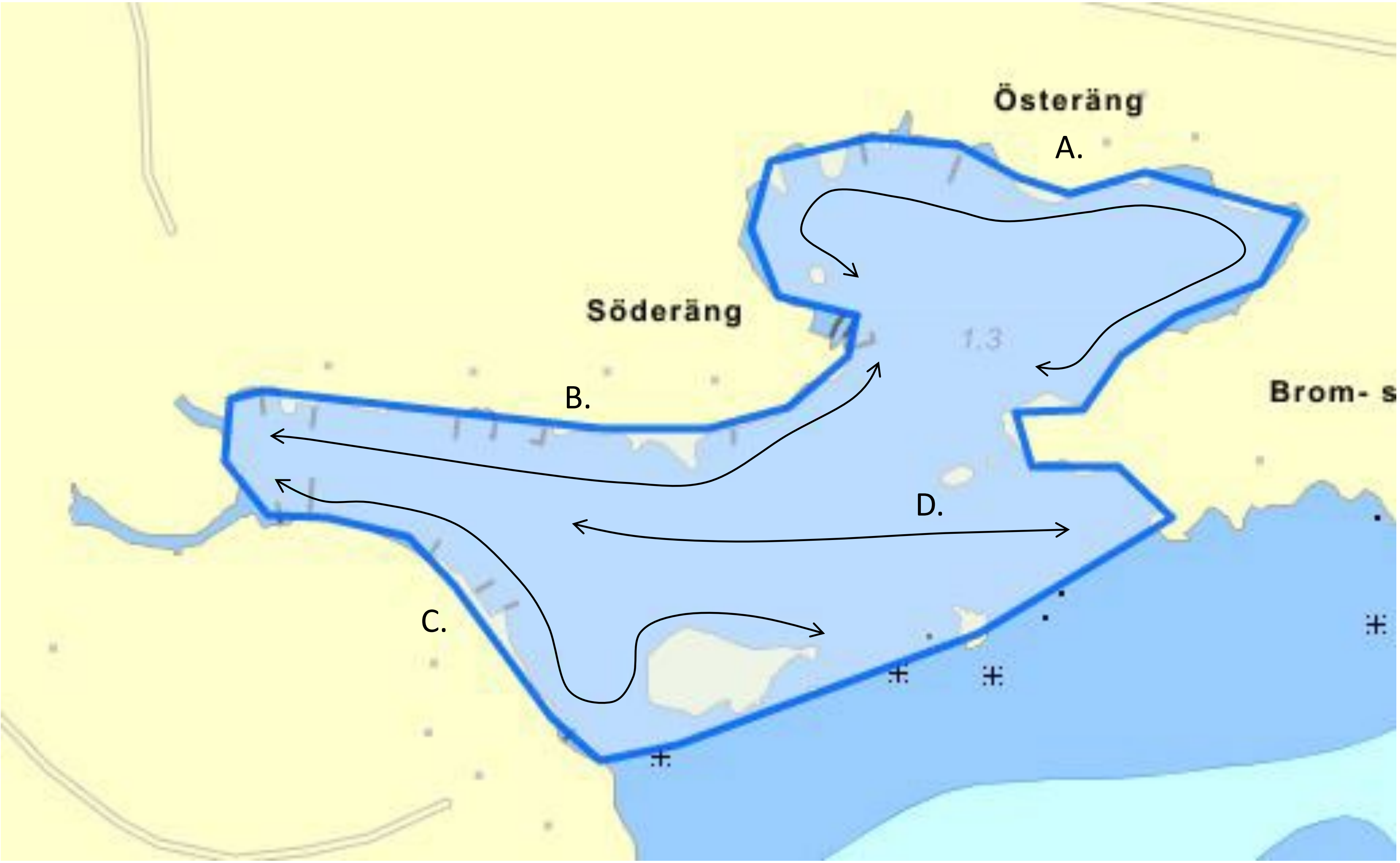

### Tomtviken/Urö
# (TV)
# 31 ha

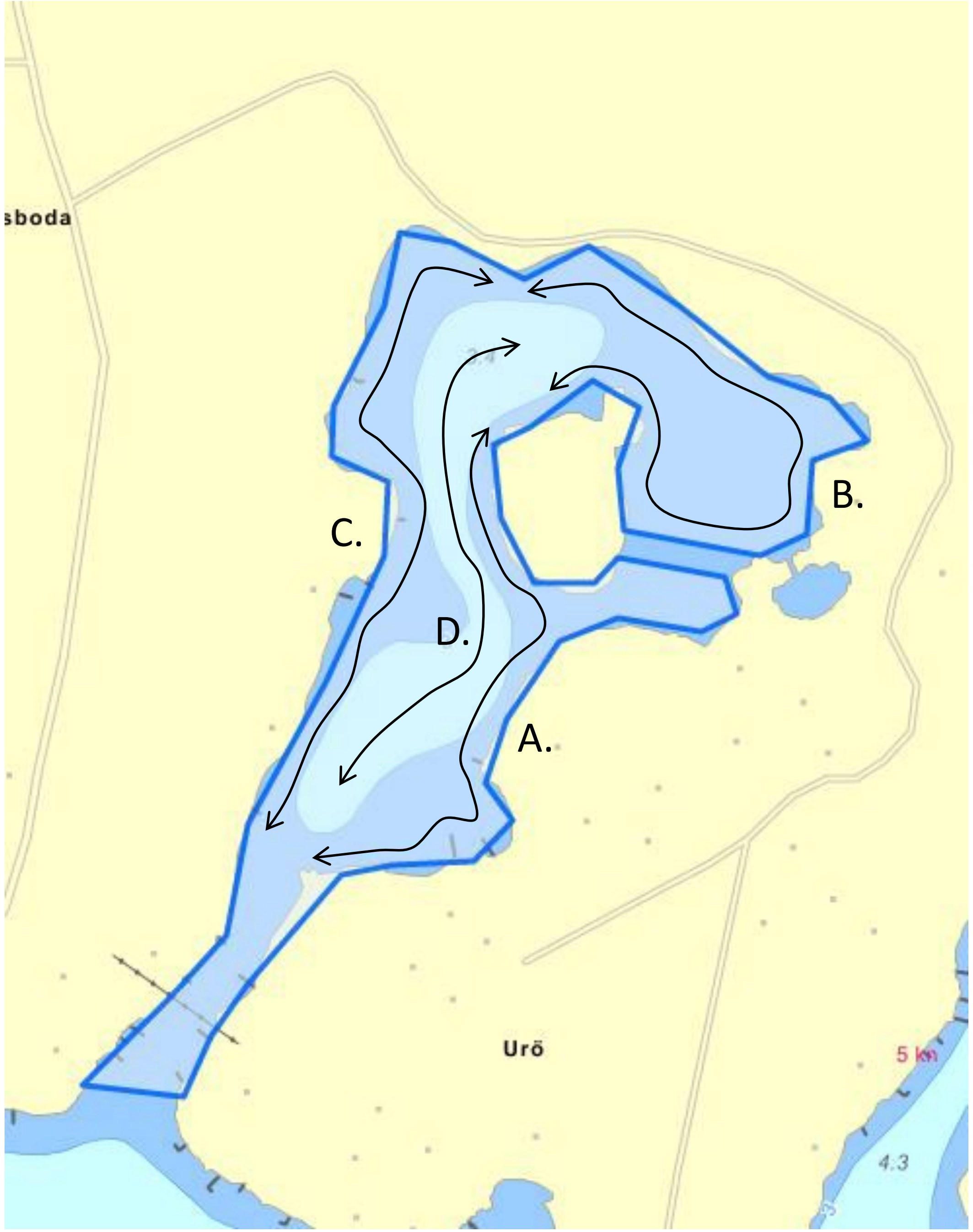

### Mulö (MÖ) 34 ha

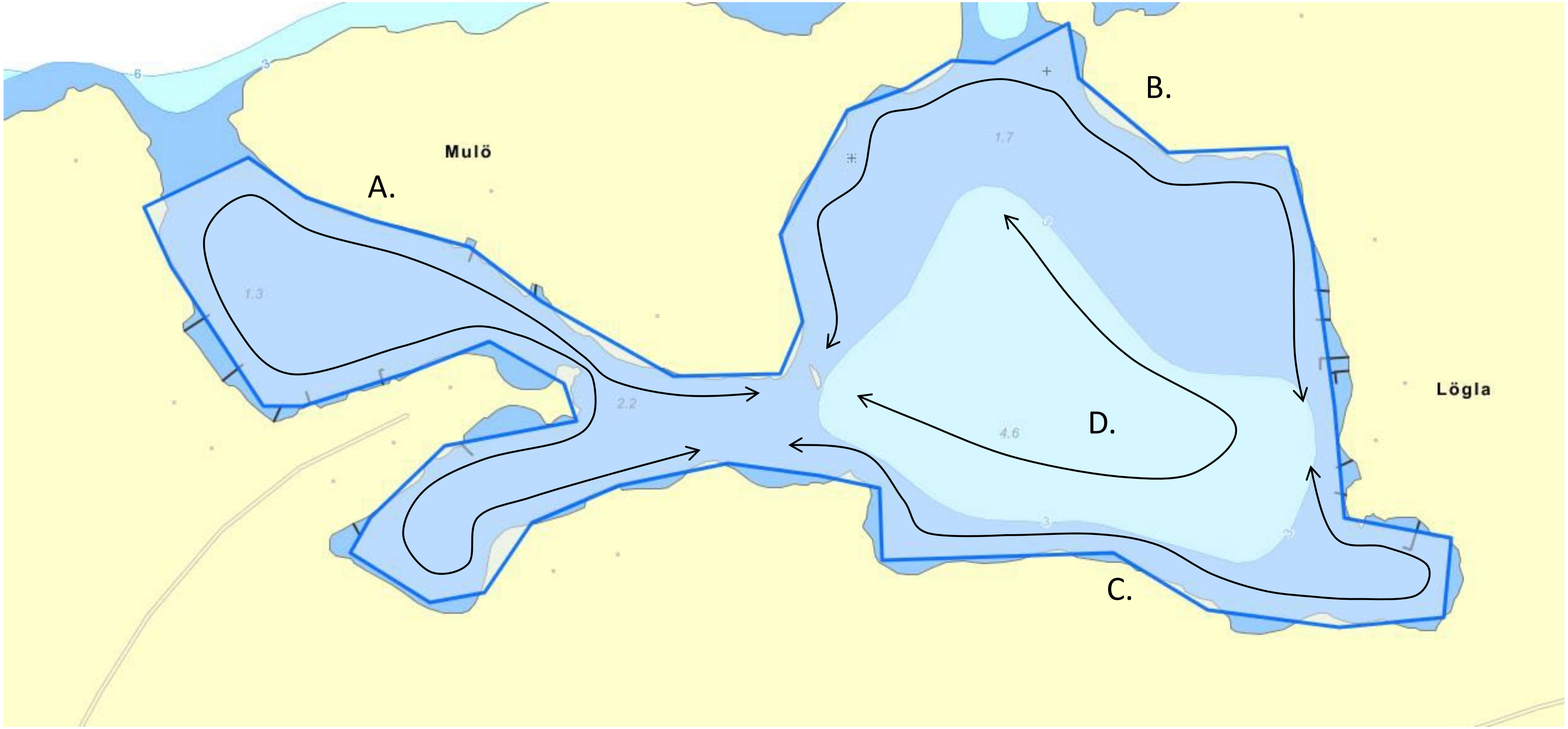
